# Metabolic and transcriptional adaptations to phagocytosis sustain microglia functionality and regenerative properties

**DOI:** 10.1101/2025.09.18.676800

**Authors:** Mar Marquez-Ropero, Jorge Valero, Xabier Cuesta-Puente, Angel Marquez-Galera, Marta Pereira-Iglesias, Alice Louail, Marco Gonzalez-Dominguez, Sol Beccari, Luzia Lopez-Murillo, Amaia San Juan, Guillermo Arrey, Irune Diaz-Aparicio, Montserrat Puigdelloses-Valcobra, Marissa L. Dubbelaar, Alberto Ribera-Ramos, Manuel Sarmiento-Soto, Rocío Ruiz, Leyre Ayerra, Amaia Vilas-Zornoza, Diana Cabrera, Sebastiaan van Liempd, Fernando García-Moreno, Klas Blomgren, Susana González-Granero, Jose M. García-Verdugo, Juan M. Falcón-Pérez, María S. Aymerich, Dolores Hambardzumyan, Iñigo Casafont, Bart Eggen, José L. Venero, Jose P. López-Atalaya, Amanda Sierra

**Affiliations:** Achucarro Basque Center for Neuroscience, Leioa, Spain; Department of Neuroscience, University of the Basque Country EHU/UPV, Leioa, Spain; Department of Cell Biology and Pathology, Universidad de Salamanca, Salamanca, Spain; Institute of Biomedical Research of Salamanca, Salamanca, Spain; Institute of Neuroscience, Molecular Neurobiology and Neuropathology Unit, Alicante, Spain; Departments of Oncological Sciences and Neurosurgery, the Tisch Cancer Institute, Icahn School of Medicine at Mount Sinai, New York, New York, USA; Department of Biomedical Sciences, University of Groningen, University Medical Center Groningen, Groningen, The Netherlands; Institute of Biomedicine of Seville, IBiS/Hospital Universitario Virgen del Rocio/CSIC/University of Seville, and Department of Biochemistry and Molecular Biology, Faculty of Pharmacy, Spain; University of Navarra, Department of Biochemistry and Genetics, Pamplona, Spain; University of Navarra, CIMA, Neuroscience Program, Pamplona, Spain; University of Navarra, CIMA, Oncohematology Program, Pamplona, Spain; Metabolomics Platform, Center for Cooperative Research in Biosciences (CIC bioGUNE), Basque Research and Technology Alliance (BRTA), 48160 Derio, Spain; Ikerbasque Foundation, 48013, Bilbao, Spain; Karolinska Institute, Department of Women’s and Children’s Health, Stockholm, Sweden; Laboratory of Comparative Neurobiology, Cavanilles Institute of Biodiversity and Evolutionary Biology, University of Valencia and CIBERNED-ISCIII, Valencia, Spain; Exosomes Laboratory and Metabolomics Platform, Center for Cooperative Research in Biosciences (CIC bioGUNE), Basque Research and Technology Alliance (BRTA), 48160 Derio, Spain; Centro de Investigación Biomédica en Red de Enfermedades Hepáticas Y Digestivas (CIBERehd), 28029 Madrid, Spain; University of Cantabria-IDIVAL, Department of Anatomy and Cell Biology, Santander, Spain; Department of Biochemistry and Molecular Biology, University of the Basque Country EHU/UPV, Leioa, Spain

**Keywords:** Phagocytosis, apoptosis, microglia, efferocytosis, metabolism, scRNA-Seq, polyamines, galectin 3, glioblastoma, irradiation

## Abstract

Phagocytosis of apoptotic cells, or efferocytosis, is a tightly regulated process that ensures tissue homeostasis and prevents mounting inflammatory responses. In the brain parenchyma, it is executed by microglia, which are encumbered by large numbers of apoptotic debris generated during development, in adult neurogenic niches, aging, and brain diseases. Emerging evidence suggest that phagocytosis is not limited to garbage disposal, but triggers adaptations in the phagocytes that may have a functional impact. To test it we developed an in vivo model of superphagocytosis induced by low cranial irradiation (LCI, 2Gy) that specifically induced apoptosis in the neurogenic niche of the adult hippocampus, synchronizing microglia in a phagocytic state within 6h and leading to full clearance by 24h. Single cell RNA sequencing and metabolomics revealed an unexpected oxidative stress in post-phagocytic microglia, accompanied by catabolic shutdown, mitochondrial remodeling, increased expression of galectin 3, and production of polyamines that led to cell death and compensatory proliferation. To test whether these changes impaired subsequent microglial phagocytosis, we used a glioblastoma model treated with sequential irradiation to induce tumor cell apoptosis. The phagocytosis efficiency of tumor-associated microglia/macrophages was comparable in the first and second apoptotic challenge, suggesting that the metabolic remodeling induced by phagocytosis was adaptive and destined to sustain their functionality. Finally, we assessed the functional impact of post-phagocytosis adaptations using galectin 3 deficient mice under LCI. We found that the recovery of the neurogenic niche after LCI strongly depended on galectin 3, demonstrating the regenerative capacity of post-phagocytic microglia. Overall, our data unveils the complexity of post-phagocytosis adaptations in microglia, underscoring their unexplored therapeutic potential in brain disorders.

## INTRODUCTION

An integral component of the braińs response to damage is apoptotic cell phagocytosis (efferocytosis), which is largely executed by microglia^1^. Microglial phagocytosis is essential to maintain tissue homeostasis, as it swiftly removes cell debris that would otherwise activate pathogen and disease-associated molecular patterns (PAMPs and DAMPS, respectively), thus preventing inflammatory responses^2^. Several transcriptional states have been related to phagocytosis in physiological and pathological conditions, including DAM (disease-associated microglia)^3^, PAM (proliferative-region associated microglia)^4^, and an interferon (IFN) I-responsive microglia^5^. However, a critical aspect that has been widely neglected is the functional impact of phagocytosis beyond debris clearing, when microglia are dealing with the consequences of cargo digestion.

Phagocytosis of apoptotic cells leads to metabolic adaptations in the phagocyte itself and reprograms its function. For example, in bone marrow-derived macrophages, phagocytosis triggers mitochondrial fission^6,7^ and decreased mitochondrial membrane potential via mitochondrial uncoupling protein 2 (UCP2)^8^, which, together with increased glycolysis^7^, ensures the continued uptake of apoptotic corpses. During *Drosophila* early development, naïve macrophages are unresponsive to tissue damage or infection but, upon apoptotic cell phagocytosis, they migrate into damaged regions and phagocytose bacterial pathogens, through the activation of the Jun kinase-signaling pathway and increased expression of the damage receptor Draper^9^. In mice, the secretome of phagocytic microglia impacts their neighbor cells and shapes adult hippocampal neurogenesis^10^, where microglia phagocytose the excess of newborn cells that undergo apoptosis^11^. However, the precise mechanism underlying the regenerative properties of post-phagocytic microglia remains to be determined.

To specifically identify the post-phagocytosis adaptations in microglia and their long-term functional impact, here we have developed an *in vivo* model of superphagocytosis induced by low cranial irradiation (LCI). LCI induced apoptosis of proliferating newborn cells in the subgranular zone (SGZ) of the hippocampus that synchronized dentate gyrus (DG) microglia in a phagocytic state at 6h. Clearance was completed by 24h, allowing us to study post-phagocytosis adaptations. Using single-cell RNA sequencing (scRNA-Seq), we discovered a time-dependent transcriptional signature induced by phagocytosis characterized by increased expression of lysosomal genes *Lgals3* and *Cd68* at 24h; and proliferative genes *Mik67* and *Top2a* at 7d. To understand the impact of these changes, we then used an in vitro model of phagocytosis and showed that post-phagocytic microglia endure oxidative stress, increased production of polyamines, and cell death in early time points (24h), and compensatory proliferation later on (7-30d). These changes suggest that phagocytosis is a stressful process for microglia, despite their nickname “professional brain phagocytes”. Importantly, these changes were adaptive, as they allowed microglia to cope with consecutive apoptotic challenges and maintain their phagocytic efficiency in a model of irradiated glioblastoma. Finally, we tested the impact of disturbing the post-phagocytosis adaptations using mice deficient for *Lgals3*, which encodes for galectin 3. After LCI, galectin 3 deficient mice showed a transient reduction in phagocytosis without affecting microglial numbers, but were unable to regenerate the neurogenic niche. Together, our data illuminates the adaptations to the metabolic overload imposed by apoptotic cargo, and underscores the regenerative properties of post-phagocytic microglia.

## METHODS

### Mice

All experiments were performed in mice expressing the enhanced green fluorescent protein, under the control of the mouse colony stimulating factor 1 receptor (fms-EGFP) (MacGreen, B6. Cg-Tg (Csf1r-EGFP)1Hume/J; Jackson Laboratory stock #018549) mice. In the fms-EGFP mice microglia constitutively express the green fluorescent protein (GFP) under the expression of c-fms gene^12,13^. Single cell RNA-Seq experiments were performed in wild type (WT) mice. *Nestin-tva* transgenic mice were used for tumor induction experiments^14,15^. All animals were housed in a climate controlled, pathogen-free facility with *ad libitum* food and water under a 12-hour light/dark cycle. All mice used were in a C57BL/6 background. All procedures conducted in Spain followed the European Directive 2010/63/EU and NIH guidelines, and were approved by the Ethics Committees of the University of the Basque Country EHU/UPV (Leioa, Spain; CEBA/205/2011, CEBA/206/2011, CEIAB/82/2011, CEIAB/105/2012). All procedures conducted in tumor-bearing mice were approved by the Institutional Animal Care and Use Committee (IACUC) of the Icahn School of Medicine at Mount Sinai (Protocol #2019-00619).

### Low cranial irradiation

For the in vivo LCI model, 2-month-old-fms-EGFP mice were anaesthetized with 2.5% avertin (10 µl/gr) and exposed to X irradiation using a model Yxlon Smart 200 tube (Yxlon International, Hamburg, Germany) at RT. Only the head of the animals was exposed to irradiation, while the rest of the body was covered with a lead blanket. The radiation was delivered as a single dose of 2 Gy at a dose rate of 0.92 Gy/min during a total time of 2 min. The source-half-depth distance was initially calculated to obtain a constant dose rate of 0.92 Gy/min. After exposure mice were sacrificed at 6h, 24h, 3d, 7d and 30d by transcardial perfusion with 30ml of PBS followed by 30ml of 4% PFA. In the double irradiation experiments, animals were sequentially exposed to a 2Gy X-ray cranial irradiation twice, 7 days apart, and were sacrificed at 6h and 24h.

### Induction of glioblastoma

DF-1 cells (ATCC, CRL-12203, RRID:CVCL_0570) were purchased and grown at 39 °C according to the manufacturers’s instructions. Injections of DF-1 cells were performed using a stereotactic fixation device (Stoelting, Wood Dale, IL). Mice used for these experiments were adults ranging from 6 to 12 weeks old, as previously utilized^16^. Mice were anaesthetized with intraperitoneal (ip) injections of ketamine (0.1 mg/g) and xylazine (0.01 mg/g). For brain injections, one microliter of 4 × 10^4^ transfected DF-1 cells in suspension was delivered using a 30-gauge needle attached to Hamilton syringe. A 1:1 mixture of cells and culture medium was used for co-injections. Locations were determined according to the coordinates in the mouse brain atlas (Franklin et al., 2019). Coordinates for SVZ injections were: AP-0mm from bregma, Lat-0.5 mm (right of midline), and depth-1.5 mm from the dural surface. Right frontal striatum coordinates were: AP-1.7 mm from Bregma, Lat-0.5 mm (left or right), and depth-2 mm from the dural surface. Mice were monitored carefully and sacrificed when they displayed endpoint symptoms. For the irradiation of GBM, tumor bearing mice were monitored and irradiated when they started to display endpoint symptoms (approximately 2 months after the injection). Mice were anaesthetized with ketamine (0.1 mg/g) and xylazine (0.01 mg/g), and exposed to X irradiation using the RS 2000 from Rad Source. Only the head of the animals was exposed to irradiation, while the rest of the body was shielded with a lead cover in a homemade apparatus. The radiation was delivered as a single dose of 10 Gy at a dose rate of 1.2 Gy/min during a total time of 8 min. After exposure, mice were sacrificed at 6h and 24h, by transcardial perfusion with 30ml of PBS followed by 30ml of 4% PFA. The tissue was posteriorly processed for immunofluorescence.

### Cell cultures

SH-SY5Y (American Type Culture Collection), a human neuroblastoma cell line was used for phagocytic assay experiments. Vampire-SH-SY5Y cells were developed through stable transfection of the SH-SY5Y cell line with tFP602 (InnoProt, P20303), and expressed red fluorescent protein as a free cytoplasmic protein. SH-SY5Y cells were grown as an adherent culture in non-coated culture flasks covered with 10-15ml of Dulbecco’s Modified Eagle Medium (DMEM, PAN-Biotech), supplemented with 10% Fetal Bovine Serum (FBS) (PAN-Biotech), 1% antibiotic/antimycotic (Gibco), and 0.5% geneticin (Fisher). When 80-90% confluency was reached, cells were trypsinized and replated at 1:4. BV2 cells (Interlab Cell Line Collection San Martino-Instituto Scientifico Tumori-Instituto Nazionale per la Ricerca sul Cancro), a line derived from raf/myc-immortalized murine neonatal microglia, was used to perform phagocytosis *in vitro* assays for the analysis of the mitochondrial network. BV2 cells were grown as an adherent culture in non-coated culture flasks covered with 10ml DMEM (PAN-Biotech), supplemented with 10% FBS (PAN-Biotech) and 1% antibiotic/antimycotic (Gibco). When confluency was reached, cells were trypsinized and replated at 1:4. For experiments, BV2 cells were counted and plated at a density of 80,000 cell/swell on poly-l-lysine-coated glass coverslips in 24-well plates (BioLite, Thermofisher). Primary microglia cultures were performed as previously described^10^. Postnatal day 0-1 (P0-P1) fms-EGFP mice pup brains were extracted and the meninges were peeled off in Hank’s balanced salt solution (HBSS, Gibco) under a binocular magnifier. The olfactory bulb and cerebellum were discarded and the rest of the brain was then mechanically homogenized by careful pipetting and enzymatically digested in an enzymatic solution (116mM NaCl, 5.4mM KCl, 26mM NaHCO_3_, 1mM NaH_2_PO_4_, 1.5mM CaCL_2_, 1mM MgSO4, 0.5mM EDTA, 25mM glucose, 1mM L-cysteine) with papain (20U/ml, Sigma), and deoxyribonuclease (DNAse; 150U/μl, Invitrogen) for 15min at 37°C. The resulting cell suspension was then filtered through a 40μm nylon cell strainer (Fisher) and transferred to a 50 ml Falcon tube quenched by 5 ml of 20% FBS(Gibco) in HBSS. Afterwards, the cell suspension was centrifuged at 200 g for 5 min, the pellet was resuspended in 1ml DMEM(PAN-Biotech) complemented with 10% FBS (PAN-Biotech),1% Antibiotic/Antimycotic (Gibco), and 5ng/ml granulocyte-macrophage colony stimulating factor (GM-CSF, Sigma), and seeded in poly-L-lysine-coated (15μl/ml, Sigma) culture flasks with a density of two brains per flask. Medium was changed the day after and then every 3–4 days. After 100% confluence (approximately 11-14 days in culture), microglia cells were harvested by shaking at 140 rpm, 37°C, 4h. DF-1 cell line (American Type Culture Collection, CRL-3586), a spontaneously immortalized fibroblast cell line was transfected with viruses and consecutively employed in brain injections to induce brain tumors. DF-1 cells were grown as an adherent culture in non-coated culture T25 flasks covered with 6ml of DMEM (ATCC), supplemented with 10% FBS (ATCC), 1% penicillin/streptomycin (Gibco) and 1% GlutaMax (Gibco). Cells were grown at 39°C according to ATCC instructions, used in early passages, and tested for mycoplasma. Cells were split every other day using trypsin and replated at 1:8. The day of the transfection cells were 30-50% confluent.

### Transfections

GFP-Mito-7 plasmid (Addgene, #57148) was transfected into BV2 cells using Lipofectamine 2000 Mito-7 corresponds to the mitochondrial targeting sequence from subunit VII of cytochrome C oxidase (Kitay et al., 2013). BV2 cells were cultured in poly-lysine-treated coverslips in a 24-well plate at a density of 50.000 cells/well. Transfection media was then changed to DMEM 10% FBS, 1% antimycotic-antibiotic.Transfections of DF-1 cells with retroviral vectors derived from avian sarcoma-leukosis virus (RCAS): RCAS-shRNA-Nf1, RCAS-hPDGFA-myc, RCAS-shRNA-p53-RFP, RCAS-shRNA-Pten-RFP for inducing mesenchymal GBM subtypes were performed using a Fugene 6 transfection kit (Roche #11814443001) according to the manufacturer’s protocol.

### In vitro phagocytosis assays

Primary microglia and BV2 cells were allowed to settle after plating for at least 24h before phagocytosis experiments. In some experiments (Seahorse, metabolomics, ROS measurement), microglia were pre-cultured in low glucose medium (2.5mM) for three days prior to the phagocytosis assay. Vampire SH-SY5Y cells were treated with 3 μM staurosporine (Sigma) for 4h at 37°C to induce apoptosis. The floating dead cell fraction was collected from the supernatant and apoptotic cells were visualized and quantified by trypan blue staining in a Neubauer chamber. Because cell membrane integrity is still maintained in early induced apoptotic cells, trypan blue unlabeled cells were considered apoptotic. Primary microglia were fed with apoptotic Vampire SH-SY5Y cells by adding them at a 1:3 proportion for Seahorse, metabolomics, and ROS measurement experiments, and the non-engulfed cells were washed out after 5h. For TEM experiments BV2 microglia were fed for 1h with apoptotic SH-SY5Y by adding the cells in a 1:1 proportion. All the analyses were performed 24h after adding the apoptotic cells.

### Seahorse extracellular flux analysis

Primary microglia were plated at a density of 100,000 cells/well in a Seahorse XFe96 cell culture microplate (Agilent) in a cell suspension of 80 µl/well to avoid cells attaching to the well’s walls. The four corner wells were not seeded and were used as background. 24h after plating, the media was changed and cells were maintained in low glucose DMEM (2.5mM) for 3 days before performing the phagocytosis assay. The analysis of the mitochondrial function in naïve and phagocytic microglia was done using the Seahorse XF Cell Mito Stress Kit (Agilent), which targets several components of the mitochondrial ETC. 1 µM Oligomycin, 1 µM FCCP, and 0.5 µM rotenone/0.5 µM antimycin A were sequentially injected in an XFe96 Sensor Cartridge (Agilent) pretreated with XF Calibrant solution (200 µl/well) overnight at 37°C in a non-CO_2_ incubator. XF Seahorse Medium (1mM pyruvate (Merck), 2mM glutamine (Sigma), 2.5 mM glucose (Sigma) and HEPES buffer 5 mM) were added in each well, including those at the corners (background correction or blank wells). The media was heated at 37°C and then the pH was adjusted to 7.4 with NaOH 1N. Cells were maintained with the supplemented medium for 1h before assay in a non-CO_2_ incubator (180 µl/ well). The analysis of the glycolytic function was performed using the Seahorse XF Glycolysis Rate Assay Kit (Agilent). 0.5 µM rotenone/0.5 µM antimycin A and 50mM 2-DG were sequentially injected in an XFe96 Sensor Cartridge pretreated with XF Calibrant solution (200 µl/well) O/N at 37°C in a non-CO_2_ incubator. DMEM 10% FBS, 1% Antimycotic/antibiotic was replaced by XF Seahorse Medium (1mM pyruvate (Merck), 2mM glutamine (Sigma), 2.5 mM glucose (Sigma), and HEPES buffer 5 mM for each well including background wells. The media was heated at 37°C and then the pH was adjusted to 7.4 with NaOH 1N. Cells were maintained in supplemented medium for 1h before assay in a non-CO_2_ incubator (180 µl/ well).

### Metabolic analysis of the tricarboxylic (TCA) and Methionine cycle metabolites

Primary microglia were plated at a density of 1 million cells/well in 6-well plates (BioLite, Thermofisher). 3 wells were seeded for each experimental condition (naïve/phagocytic) and for each metabolic pathway (TCA/Methionine cycle). 24h after plating, the culture media changed and cells were maintained in low glucose concentration (2.5mM) for 3 days before performing the phagocytosis assay. 24h after adding apoptotic cells all the wells were washed 3-times with ice-cold PBS 1X, the plate was sealed with Parafilm (Bemis, Parafilm) and then frozen at −80°C. For the extraction of metabolites of the TCA cycle 500 µL ice-cold methanol/water (50/50 %v/v) was added to the wells of the culture plates (6-wells-plate). The plates were left on dry ice for 15 minutes. Subsequently 400 µL of the homogenate plus 400 µL of chloroform was transferred to a new aliquot and shaken at 1400 rpm for 60 minutes at 4 °C. Next, the aliquots were centrifuged for 30 minutes at 13000 rpm at 4 °C. The organic phase was separated from the aqueous phase. From the aqueous phase 250µL was transferred to a fresh aliquot and placed at −80 °C for 20 minutes. 150 µL of supernatants were evaporated with a SpeedVac in approximately 1.5h. The resulting pellets were each resuspended in 150 µL water/acetonitrile (MeCN)(40/60v/v/%). Samples were measured with a UPLC system (Acquity, Waters Inc., Manchester, UK) coupled to a Time-of-Flight mass spectrometer (ToF MS, SYNAPT G2, Waters Inc.). A 2.1 x 100 mm, 1.7 µm BEH amide column (Waters Inc.), thermostated at 40 °C, was used to separate the analytes before entering the MS. Mobile phase solvent A (aqueous phase) consisted of 99.5% water and 0.5% FA while solvent B (organic phase) consisted of 4.5% water, 95% MeCN and 0.5% formic acid (FA). In order to obtain a good separation of the analytes the following gradient was used: from 10% A to 99.9% A in 2.6 minutes in curved gradient (#9, as defined by Waters), constant at 99.9% A for 1.6 minutes, back to 10% A in 0.3 minutes. The flow rate was 0.250 mL/min and the injection volume was 4 µL. After every 8 injections sample was injected. The MS was operated in negative electrospray ionization in full scan mode. The cone voltage was 25 V and capillary voltage was 250 V. Source temperature was set to 120 °C and capillary temperature to 450 °C. The flow of the cone and desolvation gas (both nitrogen) were set to 5 L/h and 600 L/h, respectively. A 2 ng/mL leucine-enkephalin solution in water/acetonitrile/formic acid (49.9/50/0.1 %v/v/v) was infused at 10 µL/min and used for a lock mass which was measured every 36 seconds for 0.5 seconds. Spectral peaks were automatically corrected for deviations in the lock mass. Extracted ion traces for relevant analytes were obtained in a 20 mDa window in their expected m/z-channels. These traces were subsequently smoothed and peak areas integrated with TargetLynx software. Signals of labelled analytes were corrected for naturally occurring isotopes. These calculated raw signals were adjusted for by median fold-change (MFC) adjustment. This is a robust adjustment factor for global variations in signal due to differences in tissue amounts, signal drift or evaporation. The MFC is based on the total amount of detected mass spectrometric features (unique retention time/mass pairs). The calculations and performance of the MFC adjustment factors were performed as previously described^17,18^. For extraction of metabolites of the Methionine cycle 500 µL ice cold methanol containing 200 mM acetic acid was added to the wells of the culture plates (6-wells-plate). The plates were left on dry ice for 15 minutes. Subsequently the extraction liquid was transferred from the wells to Eppendorf tubes and centrifuged at 3750 rpm at 4 °C during 30 minutes. 250 µL of the chilled supernatants were evaporated with a SpeedVac in approximately 1.5 h. The resulting pellets were resuspended in 150 µL water/acetonitrile (40/60 v/v/). Samples were measured with an ultra-performance liquid chromatographic system (Acquity, Waters Inc., Manchester, UK) coupled to a Time-of-Flight mass spectrometer (SYNAPT G2, Waters Inc.). A 2.1 x 100 mm, 1.7 µm BEH amide column (Waters Inc.), thermostated at 40 °C, was used to separate the analytes before entering the MS. Mobile phase solvent A (aqueous phase) consisted of 99.5% water, 0.5% FA and 20 mM ammonium formate while solvent B (organic phase) consisted of 29.5% water, 70% ACN, 0.5% FA and 1 mM ammonium formate. In order to obtain a good separation of the analytes the following gradient was used: from 5% A to 50% A in 2.4 minutes in curved gradient (#8, as defined by Waters), from 50% A to 99.9% A in 0.2 minutes constant at 99.9% A for 1.2 minutes, back to 5% A in 0.2 minutes. The flow rate was 0.250 mL/min and the injection volume was 4 µL. All samples were injected randomly. After every 8 injections QC sample (pooled samples) was injected. The MS was operated in positive electrospray ionization mode in full scan (50 Da to 1200 Da). The cone voltage was 25 V and capillary voltage was 250 V. Source temperature was set to 120 °C and capillary temperature to 450 °C. The flow of the cone and desolvation gas (both nitrogen) were set to 5 L/h and 600 L/h, respectively. A 2 ng/mL leucine-enkephalin solution in water/acetonitrile/formic acid (49.9/50/0.1 %v/v/v) was infused at 10 µL/min and used for a lock mass which was measured each 36 seconds for 0.5 seconds. Spectral peaks were automatically corrected for deviations in the lock mass.

### ROS analysis

Primary microglia were plated at a density of 40,000 cells/well on poly-L-lysine-coated 96-well plates (Thermofisher) and the cells were allowed to rest and settle for at least 24h before phagocytosis experiments. 1h before the measurement the culture medium was replaced and cells were incubated with CellROX (5 µM, Invitrogen) and the nuclear dye NucBlue (8µl/well, Invitrogen) added to phenol-free DMEM 10% FBS, 1% antimycotic-antibiotic, 2.5mM glucose, 1mM pyruvate in the incubator of the CellInsight CX7 Pro high-content screening system (ThermoFisher) at 37°C and 5% CO2. After incubation images were taken at 644/665 nm using a 40x (0.6 NA) Olympus objective for a maximum time of 2 hours.

### Immunofluorescence and immunohistochemistry

Primary microglial cultures were fixed for 10 min in 4% PFA and then transferred to PBS. Fluorescent immunostaining was carried out following standard procedures^19^. Coverslips with primary microglial cultures were blocked in 0.1% Triton X-100, 0.5% BSA in PBS for 30 min at RT. The cells were then incubated with primary antibodies (**Table 1**) in blocking solution (0.1% Triton X 100, 0.5% BSA in PBS) overnight at RT, rinsed in PBS and incubated in the secondary antibodies containing DAPI (5 mg/ml) in the permeabilization solution for 1h at RT. After washing with PBS, primary cultures were mounted on glass slides with Dako Fluorescent Mounting Medium (Agilent). Mice were transcardially perfused with 30ml of PBS followed by 30ml of 4% PFA. The brains were postfixed with the same fixative for 4h at RT, then washed in PBS and kept at 4°C. Six series of 50μm-thick sagittal sections of mouse brains were cut using a Leica VT 1200S vibrating blade microtome (Leica Microsystems GmbH, Wetzlar, Germany). Fluorescent immunostaining was carried out following standard procedures^19^. Free-floating vibratome sections were blocked in permeabilization solution (0.3% Triton-X100, 0.5% BSA in PBS; all from Sigma) for 2 hr at RT, and then incubated overnight with the primary antibodies (**Table 1**) diluted in the permeabilization solution at 4°C. For BrdU (bromodeoxyuridine), activated caspase 3 and Iba1 labeling an antigen retrieval procedure was performed by incubating in 2M HCl for 20min at 37°C and then washing with 0.1M sodium tetraborate for 15min at RT (3 washes of 5min each) prior to the blockade of the sections. After overnight incubation with primary antibodies, brain sections were thoroughly washed with 0.3% Triton in PBS. Next, the sections were incubated with fluorochrome-conjugated secondary antibodies and DAPI (5mg/ml; Sigma) diluted in the permeabilization solution for 2h at RT. After washing with PBS, the sections were mounted on glass slides with DakoCytomation Fluorescent Mounting Medium (Agilent). For immunohistochemistry of activated-caspase 3 in glioblastoma, mice were transcardially perfused with 30ml of PBS followed by 30ml of 4% PFA. 24h post-fixation, brains were embedded in 30% sucrose-DEPC treated until the brains sank and then were frozen at −80°C in O.C.T. Compound (Cell Path). 20μm sections were cut onto Superfrost Plus slides using a LEICA CM1950 Cryostat (Leica) and kept at −20°C until use. The tissue was air-dried and antigen retrieval was performed by immersing the slides in a jar containing antigen retrieval buffer pH 6 (Vector labs) diluted in Milli-Q H_2_O. The jar was inserted into a container with boiling water and the samples were maintained at 93°C for 15-25min. Samples were put back to RT in an histology box and blocked in permeabilization solution (0.3% Triton X-100 (Sigma) and 5% normal goat serum in PBS) for 1h at RT followed by 3% H_2_O_2_ incubation for 30min. Slides were incubated in caspase 3 primary antibody (prepared in PBS) at 4°C overnight. After overnight incubation with primary antibodies, slides were washed with 0.3% triton in PBS. Next, the sections were incubated with biotinylated goat anti-rabbit secondary antibody (prepared in PBS) for 1h at RT. Secondary antibody was washed with 0.3% triton in PBS. Biotin labeling was developed by incubating the samples with ABC solution (Avidin Biotin Complex Elite Kit; Vector labs) following manufactures instructions, followed by incubation with 2% 3,3′-Diaminobenzidine tetrahydrochloride (DAB) (Sigma) diluted in 1.5% buffer stock solution, 1.5% H_2_O_2_ Samples were washed with 0.3% Triton in PBS. Nuclei staining was developed by 1min incubation in hematoxylin solution (Sigma). After washing with PBS, the sections were mounted on glass slides with 100% glycerol (Sigma).

**Table 1.**
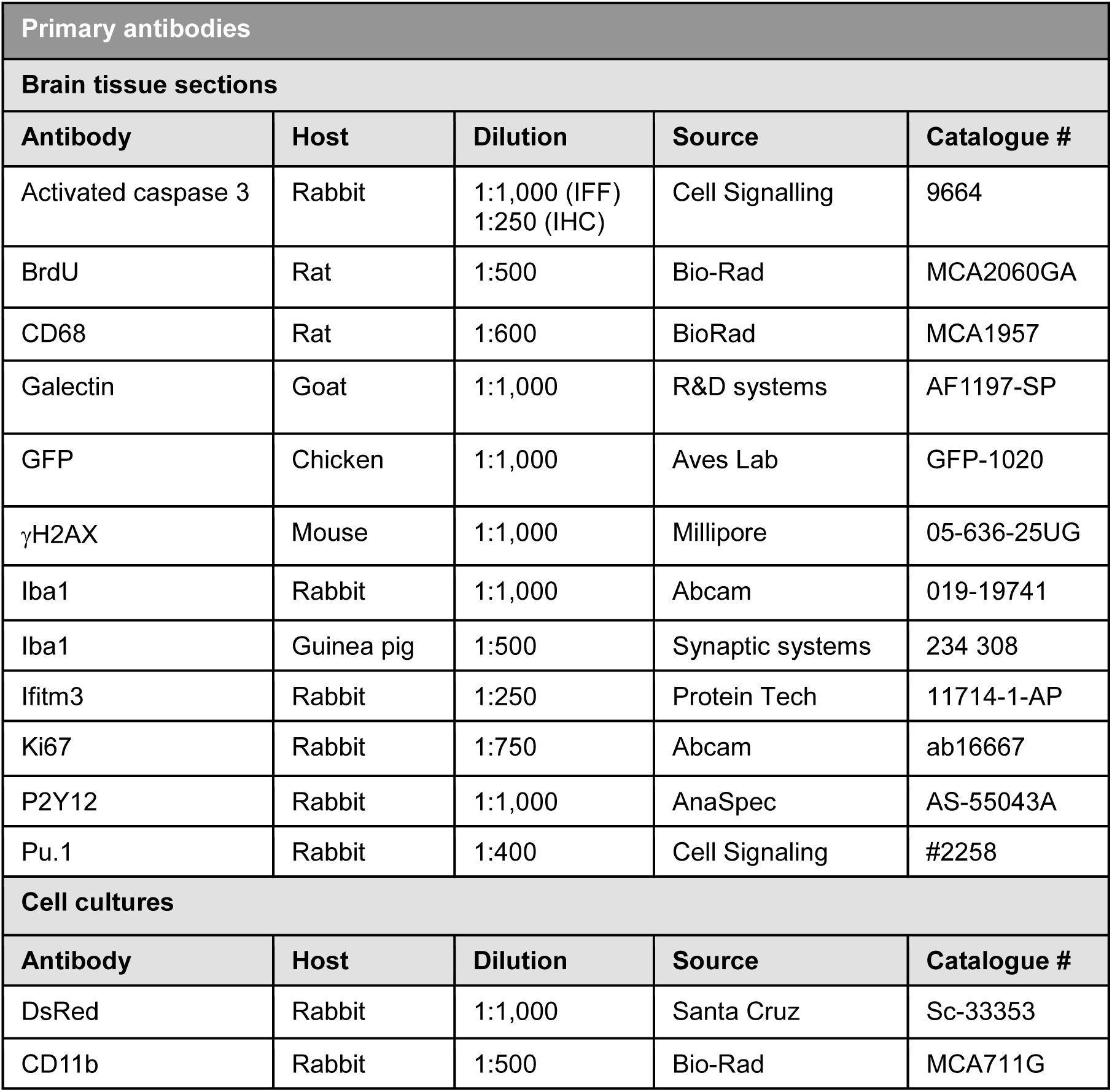
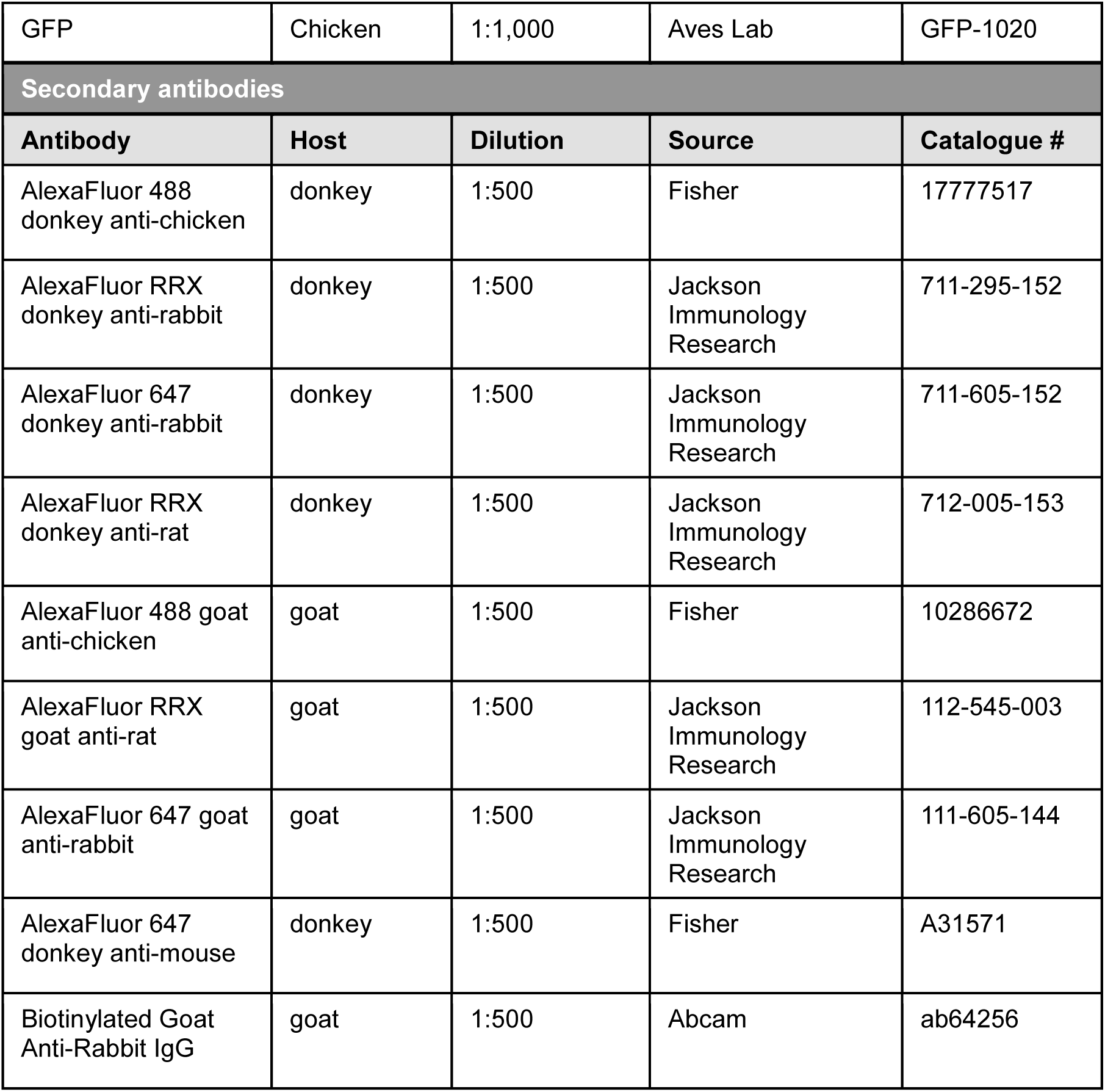
List of antibodies used.

### RNAScope

To detect mRNA, RNAscope was performed on 4% PFA-DEPC treated fixed brains. 24h post-fixation, brains were embedded in 30% sucrose-DEPC treated until sinking and frozen at −80°C in O.C.T. Compound (Cell Path). 20μm sections were cut onto Superfrost Plus slides using a LEICA CM1950 Cryostat (Leica) and dried at - 20°C for 2h and kept at −80°C. RNAscope was carried out according to manufacturer’s instructions with the following catalogue probes: Mm-*Ifit3* (ACDBio, 508251) and Mm-*Ms4a7* (ACDBio, 314601). The probe *Apoe* (ACDBio, 313271) was used as a positive internal control. For the development of the fluorescent signal the fluorophore OPAL 570 dye (Akoya) was used. RNAscope was followed by immunofluorescence with anti-P2Y12.

### Confocal imaging, image processing, and quantification

Fluorescence immunostaining images were collected using a SP8 laser scanning microscope (Leica) or a Stellaris confocal microscope (Leica) using a 40X oil-immersion objective and a z-step of 0.7 μm for phagocytosis analysis, and a 63X objective and a z-step of 0.3μm for mitochondrial morphology analysis. All images were saved in a Tiff format and used for further analysis in ImageJ. For the analysis of cells in cultures, 4 random images per coverslip were acquired. For mouse tissue sections, at least 3 15-20 μm-thick z-stacks located at random positions containing the DG and CA were collected per hippocampal section, and a minimum of 6 sections per series were analyzed. Quantitative analysis of apoptosis and phagocytosis, was performed using unbiased stereology methods as previously described^11^.

The number of apoptotic cells, phagocytosed apoptotic cells and the density of microglial cells were quantified using ImageJ image analysis software. The volumetric density was estimated using the DG volume contained in the z-stack (by multiplying the thickness of the stack by the area of the DG at the center of the stack). The estimation per hippocampus was only performed in brain tissue sections. For that purpose, the estimation per volume was multiplied by the volume of the septal hippocampus (spanning from −1mm to −2.5 in the AP axes, from Bregma), which was calculated by taking pictures of the last 6-7 brain slices cut with the vibratome, where the septal hippocampus is contained, with an epifluorescent microscope (Zeiss Axiovert) at 20X. To calculate the phagocytosis parameters, the following formulas were used^11,19^:

<u>Ph Index:</u> Proportion of apoptotic cells engulfed by microglia. Phagocytosed apoptotic cells (apo^Ph^); total apoptotic cells (apo^tot^).

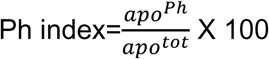

<u>Ph capacity:</u> proportion of microglia with one or more phagocytic pouches, each containing one apoptotic cell (Sierra et al., 2010). Microglia (mg); microglia with one or more phagocytic pouches (Phn).

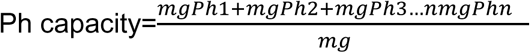

Clearance time: Ratio of apoptotic cells that disappear between 24h and 6h after LCI multiplied by the time (18h), multiplied by the Ph capacity (to account for the fact that microglia phagocytose several times at a time).

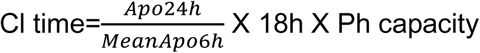

Mitochondrial analysis was performed using image analysis software ImageJ and Huygens^20^. Mitochondrial images were deconvoluted with Huygens (Scientific Volume Imaging) by using the Classic Maximum Likelihood Estimation algorithm (CMLE). Deconvolution parameters were established as 40 maximum iterations, signal-to-noise ratio 20, 0.05 of quality threshold, and optimized iteration mode. The analysis carried out with ImageJ (https://fiji.sc/) started with background removal by using the FIJI “subtract background” tool. Additionally, cell-specific background was eliminated by subtracting the cell nucleus intensity. Images were then converted to binary (black and white) by thresholding, where a foreground pixel is assigned the maximum value (255) and background pixels are assigned the minimum value (0). The “skeletonize” tool of FIJI allowed us to define the mitochondrial features of the original image (“skeleton”) and it was analyzed using the plugin Analyze Skeleton 2D/3D^21^. This plugin classified the spatial distribution of each skeleton pixel to measure the length of each branch and the number of them. Two types of mitochondrial structures were distinguished: individual mitochondria (unbranched) and mitochondrial networks (branched). Individual mitochondria were defined as those objects in the image that did not contain junction pixels and were comprised of 1 or 0 branches (defined as a single point). Mitochondrial networks were defined as those objects that contained at least 1 junction pixel and were comprised of more than 1 branch. Based on this categorization, several parameters were calculated: the number of mitochondria and their length per cell; the number of branches per cell; and the number of junctions between branches per cell. The analysis with Huygens worked with deconvoluted images. First, out of focus and noisy stacks were removed with “crop” option. Then the image was opened with the “object analyzer” tool and the values from the maximum intensity of the original image and the maximum intensity of the nucleus were selected to establish a threshold. Then the “watershed segmentation” option was selected and a sigma value of 0.2 was chosen. Objects smaller than 0.06µm^3^ were removed by applying a 38-voxel equivalent size filter, as they were not considered mitochondria. Finally, the following data from the mitochondrial objects resulting from the processing was extracted: number of mitochondrial objects, total and mean volume of objects, total and mean length of objects, and mean lateral width, axial width, roughness, sphericity, and surface of objects.

For the analysis of DNA damage, γH2AX nuclear staining was analyzed within nuclear DAPI and Pu.1 staining. 2-3 images from 6-7 brain slices were analyzed using a homemade ImageJ Macro. The Macro extracted the values for the total volume of nuclei (DAPI staining), the total volume of microglia nuclei (Pu staining) and the total volume of double-strand breaks (DSB) (γH2AX staining). Finally, the values of the % volume occupied by γH2AX in both DAPI and Pu.1 were extracted as indicators of overall nuclear damage and microglia damage, respectively. The number of γH2AX+ cells, which represented those cells that had their entire nucleus occupied by γH2AX staining, was manually quantified and finally the number of γH2AX/mm^3^ cells was extracted.

To quantify monocyte infiltration, we analyzed 2-3 images from 6-7 brain slices. Manual counting was carried out for both microglia cells (fms-EGFP^+^/P2Y12^+^) and monocytes/macrophages (fms-EGFP^+^/P2Y12^-^). Subsequently, the percentage of fms-EGFP^+^ cells identified as microglia and monocytes/macrophages was determined.

For the analysis of *Ms4a7* and *Ifit3* probes using RNAScope, 2-3 images from 6-7 brain slices were analyzed, with an average of 38 total cells analyzed per experimental condition. The labeling of *Ms4a7* and *Ifit3* probes was visually inspected and microglia were classified as *Ms4a7* ^+^ or *Ms4a7* ^-^ and I*fit3*^+^ or *Ifit3* ^-^. Subsequently, total microglia were manually counted and the % of labeled cells was calculated. In all RNAscope procedure reactions, both positive (*Apoe*) and negative (*Dapb*) controls were included.

The assessment of CD68 and Gal3 expression using immunofluorescence was conducted using at least 3 images from 6-7 brain slices. The labeling pattern differed between CD68 and Gal3. While Gal3 labeling was distributed throughout the entire cell, CD68 labeling was clustered around the nucleus and formed small accumulations along the microglial processes. Therefore, for the analysis of CD68, a threshold was established to classify microglia as CD68^+.^ This allowed us to distinguish microglia with minimal CD68 labeling, which occupied less than 5% of the microglial area. In contrast, other cells showed more evident and clustered CD68 labeling in the aforementioned regions, representing more than 5% of the microglial area occupied by CD68. For CD68 labeling, cells were considered CD68^+^ microglia if they exhibited >5% of their cytoplasm labeled with CD68, as measured by a homemade volume colocalization macro. Once CD68^+^ microglia were identified, the percentage of CD68^+^ microglia relative to the total number of microglia was determined. For Gal3 labeling, microglia expressing Gal3 were classified as Gal3^+^ microglia, and the percentage among all microglia was calculated. Additionally, Gal3^+^ microglia were classified based on low (intensity values 0-85), medium (intensity values 85-170), and high (intensity values 170-255).

### Transmission electron microscopy

Three million primary microglial cells were cultured in poly-L-lysine coated 60mm diameter plates (Nunc, Thermofisher) and the cells were allowed to rest and settle for at least 24h before phagocytosis experiments, adding apoptotic cells at a 1:1 ratio. 24h later, the cells were rinsed in PBS and pre-fixed as an adherent cell monolayer using 0.5% glutaraldehyde solution in Sörensen buffer (SB) 0.1M pH=7.4 (10 mins, RT). After scraping the pre-fixed cells, primary microglia were centrifuged (800g, 5 mins, RT) to form a pellet and fixed in 2% glutaraldehyde solution in SB (overnight, 4°C). Primary microglia were then rinsed with 4% sucrose in SB and post-fixed with 1% osmium tetroxide in SB (1h, 4°C, darkness). After rinsing, primary microglia were dehydrated in a growing concentration series of acetone. After dehydration, primary microglia samples were embedded in epoxy resin EPON Polarbed 812 (Electron Microscopy Sciences). Semi-thin sections (1μm thick) were stained with toluidine blue to identify the regions of interest. Ultra-thin sections were cut using a LEICA EM UC7 ultramicrotome and contrasted with uranyl-acetate and lead citrate. The ultrastructural analysis was done with a Transmission Electron Microscope JeoJEM 1400 Plus at 100 kVs equipped with a SCMOS digital camera. Image analysis was performed using ImageJ. Mitochondria were identified manually, and their perimeter was selected to generate regions of interest (ROIs) in each image. The area of the cellular cytoplasm (µm²) was also identified and selected manually to generate a ROI in each image. Subsequently, the following information about the mitochondrial ROIs was extracted: area, density, perimeter, circularity, ferret diameter, roughness, and solidity. Finally, the quantitative data from ROIs of the images belonging to the same cell were grouped and density was calculated dividing the number of mitochondria present in the cell by its cytoplasm area (µm²). The percentage of cytoplasm area occupied by mitochondria per cell was calculated by summing up the areas of individual mitochondria and relating it to the cellular cytoplasmic area, which was considered 100%.

In tissue sections, a pre-embedding staining protocol was used. 200 μm hippocampal sections were incubated with primary antibody rabbit anti-P2Y12 (1:500, AnaSpec) and colloidal gold-conjugated secondary antibody goat anti-Rabbit (1:50; UltraSmall; Aurion, Wageningen) as described previously ^22^. The sections were post-fixed with 1% osmium tetroxide with 7% glucose for 30 min, rinsed, dehydrated and embedded in araldite (Durcupan; Fluka, Buchs). Semithin sections (1.5 μm) were cut with an Ultracut UC-6 (Leica, Heidelberg, Germany), mounted on gelatin-coated slides, and stained with 1% toluidine blue, and examined under a light microscope (Eclipse E200, Nikon) to identify P2Y12^+^ cells. Consecutive semithin sections with P2Y12^+^ cells were re-embedded, and ultrathin sections (70 nm) were cut, stained with lead citrate (Reynolds solution) and examined under a transmission electron microscope (Tecnai Spirit G2; FEI, Eindhoven, The Netherlands). Images were acquired using Radius software (Version 2.1) with a XAROSA digital camera (EMSIS GmbH, Münster, Germany). For the ultrastructural analysis of microglia, we measured: the cytoplasmic area, the number and area occupied by lysosomes, the number and area occupied by mitochondria and the area of phagosomes. We differentiated between lysosomes or lysosomal bodies and phagosomes (containing dead cells).

### Single cell RNA sequencing

LCI-treated mice were euthanized at 24 hours and 7 days post-tratment using a terminal dose of Avertin (20 µl/gr) and perfused with PBS 1X. The hippocampi were dissected using a scalpel, and 6 hippocampi per group (from 3 animals) were pooled in a 2ml tube containing a dissociation solution with a mixture of DNase (50ug/ml) and papain (2mg/ml) dissolved in DPBS (HyClone). EDTA (10μl/tube) was added, and tubes were incubated on a Stuart S83 Rotator for shaking at 11 rpm at 37°C for 30 minutes. All subsequent stepsof the procedure were performed at 4°C, unless stated otherwise. Subsequently, the samples were homogenized using a Pasteur pipette. The resulting cell suspension was filtered through a 70 µm cell strainer (BD Bioscience) and transferred to a 15 ml tube containing1 ml DPBS. An additional 1 mL DPBS was added, and the suspension was centrifuged at 300 g for 10 min. After centrifugation, the supernatant was discarded, and the pellet was resuspended in1 ml of Percoll 25% (Cytiva) for myelin removal. The suspension was transferred to a 1.5 ml tube and centrifuged at 1,000 g (full acceleration, full brake) at room temperature for 10 minutes. The myelin-containing supernatant was discarded, and the cell pellet was resuspended in 90μl of cytometry buffer with 10μl of anti-CD11b microbead solution (Miltenyi Biotec) and incubated for 15 minutes at 4°C. After adding 1 ml of cytometry buffer, the suspension was centrifuged at 300 g for 10 min. The supernatant was removed, and cells were resuspended in 500µl cytometry buffer in a 15ml Falcon tube for magnetic separation of CD11b^+^ cells using AutoMACS (Miltenyi Biotec, “posseld” and “rise” settings). Isolated CD11b^+^ cells were resuspended at a concentration of 1,200 cells/µl for downstream processing. Cells were encapsulated using the Chromium Next GEM Chip (10X Genomics), and libraries were generated using the Chromium Controller and Single Cell 3’ Gene Expression Reagent Kits v3.1 according to the manufacturer’s protocol (10x Genomics). Library quality and quantity were assesed using a High Sensitivity DNA Kit (Agilent Technologies) on a BioAnalyzer 2100 (Agilent Technologies).

### Bioinformatic analysis of scRNA-seq

Four 10x-barcoded sequencing libraries were generated using the 10x Single Cell 3’ v3 Reagent Kits (10x Genomics) and sequenced to an average depth of approximately 300 million reads per sample (control 24h: 316,580,458; control 7d: 297,113,513; LCI 24h: 305,341,402; LCI 7d: 338,536,344). Raw FASTQ libraries were aligned to the mm10-2020-A mouse reference genome and processed using CellRanger Count (10X Genomics, Cell Ranger v5.0.1) with default parameters, yielding 15,745, 16,704, 14,295, and 18,601 cells for the respective samples. Sequencing depth was between 17,787 and 21,360 read pairs per cell, with sequencing saturation between 44.4% and 52.2%. Multiplet artifacts were removed using Scrublet v0.2.3 python package^23^ with default parameters, resulting in 15,397, 15,949, 13,813, and 18,069 cells, respectively. All subsequent single cell transcriptomics analyses were performed using R Statistical Software (v4.0.5) and the Seurat package^24^ (v4.1.1).

For quality control, genes expressed in fewer than 3 cells were removed. Cells with unique feature counts (detected genes) above 4,500 or below than 1,300, or with mitochondrial gene counts exceeding 5%, were filtered out. After filtering, 45,249 high-quality cells remained (10,955 cells in control 24h, 11,801 in control 7d, 9,281 in LCI 24h, and 13,212 in LCI 7d) for downstream analyses. Unique molecular identyfier (UMI) count data were normalized using a regularized negative binomial regression with SCTransform (v0.3.3), applying theflags vst.flavor=“v2” to invoke the v2 regularization and method=“glmGamPoi” to fit a Gamma-Poisson Generalized Linear Model which improves the speed of the learning procedure. Mitochondrial content was regressed out using a second non-regularized linear regression step. Next, to ensure that the sctransform residuals for the top variable genes across all samples are computed, we used the functions SelectIntegrationFeatures with nfeatures=3000 (getting the top 3,000 highly variable genes (HVGs) across all the samples) and PrepSCTIntegration (recomputing the residuals for missing values across the previous 3,000 HVGs using the stored model parameters). Then, the four Seurat objects were merged into a single object containing all 45,249 high quality cells with all the needed values calculated in the SCT assay, and the SCT assay set as the default assay for the downstream analysis. Principal Component Analysis (PCA) was performed on top 3,000 HVGs to reduce dimensionality. Non-linear dimensional embedding was performed using Uniform Manifold Approximation and Projection (UMAP) using the 30 most significant dimensions of the PCA. A Shared Nearest-Neighbor (SNN) graph was constructed using FindNeighbors function on the 30 most significant principal components (PCs), and cell clusters were identified using FindClusters. Differential expression analysis was conducted using Wilcoxon rank-sum test implemented in Seurat (FindMarkers, with default parameters) after running PrepSCTFindMarkers on the SCT assay.

Microglial identity was confirmed using a gene signature-based module enrichment score (AddModuleScore with default parameters)^25^. Clusters with low microglia marker scores, including macrophages and non-myeloid cells, were excluded. Microglia with a reactive profile (high score for micro/myeloid shared activation gene list^25^) and present in all the samples, likely induced by enzymatic dissociation, were also removed. UMAP was recomputed using the previous 30 most significant PCs on the remaining 32,591 high quality cells (8,453 cells in Ctrl24h, 8,961 in Ctrl7d, 6,282 in Irr24h, and 8,895 in Irr7d). PrepSCTFindMarkers function was used prior to differential testing on the SCT assay, and differential expression analysis was performed using FindAllMarkers (logfc.threshold=0.1, min.pct=0.25) to identify enriched genes in each microglial subpopulation compared to the remaining microglia. LCI-induced differentially expressed genes for each microglia subpopulation were identified using FindMarkers (logfc.threshold=0.1).

### Weighted gene co-expression network analysis (WGCNA)

gene arrays comparing cultured primary naïve and phagocytic microglia (2h, 24h) were originally published in ^10^. We performed WGCNA to identify modules of highly co-expressed genes^26^. Filtering was performed on the raw datasets of the array, and a cut-off was determined by the 95 quantile of the controls, with at least four samples having an expression higher than this cut-off before gene expression was analyzed using Ingenuity Pathway Analysis (QIAGEN; https://digitalinsights.qiagen.com/products-overview/discovery-insights-portfolio/analysis-and-visualization/qiagen-ipa/). Then, different molecular networks were generated according to biological or molecular functions, which include canonical pathways, upstream regulatory analysis, and disease-based functional networks.

### Statistical analysis

SigmaPlot (San Jose, CA, USA) and GraphPad Prism (San Diego, CA, USA) were used to perform the statistical analysis. Data was tested for normality and homoscedasticity. When the data did not comply with these assumptions, a logarithmic transformation (Log10, Log10+1, or Ln) or a square root was performed, and the data was analyzed using parametric tests. Two-sample experiments were analyzed by Student’s t-test and more than two-sample experiments with one-way or two-way ANOVA. In case that homoscedasticity or normality were not achieved with a logarithmic transformation, data were analyzed using a Kruskal-Wallis ranks test, followed Dunn method as a posthoc test. Two sample non-parametric data was analyzed using the Mann Whittney U test. Only p < 0.05 is reported to be significant.

### Data and code availiability

RNA-seq data have been deposited at the NCBI GEO database as GEO: GSE307614 and are publicly available as of the date of publication. This paper does not report original code. Any additional information to reanalyze the data reported in this paper is available from the lead contact upon request.

## RESULTS

### Low cranial irradiation (LCI) induces microglial superphagocytosis in the hippocampal DG

We developed and validated an *in vivo* model of superphagocytosis that allowed us to study post-phagocytic events in microglia up to 30d (**Fig. 1**), overcoming the limitation of having sparse phagocytic microglia due to low levels of apoptosis that are found in neurogenic niches, the only regions where physiological apoptosis occurs in the adult brain^10,11^. We used a low cranial irradiation (LCI) paradigm by administering 2Gy to the skull, a dose well-below the 8Gy commonly used in tumor irradiation studies in mice^27^. In the neurogenic niche of the hippocampus, apoptotic cells labeled with activated caspase 3 were exclusively located in the subgranular zone (SGZ) of the dentate gyrus (DG) 6h after LCI (**Fig. 1A, B**), suggesting that it specifically induced apoptosis in the proliferating neuroprogenitors residing in the SGZ. Apoptotic cells were no longer detectable by 24h (**Fig. 1C, D**), indicating effective microglial removal and an average clearance time of 41.0±7.5min per apoptotic cell. Indeed, the vast majority of apoptotic cells were engulfed by microglia, visualized in fms-EGFP mice (**Fig. 1E**), as SGZ microglia displayed multiple pouches containing apoptotic cells 6h after LCI (**Fig. 1F-H**). Distal microglia (from the molecular layer of the DG) were also recruited to engage in phagocytosis (**Fig. 1F**). Thus, DG microglia were synchronized in phagocytosis 6h after LCI, and became post-phagocytic afterwards.

**Figure 1.**
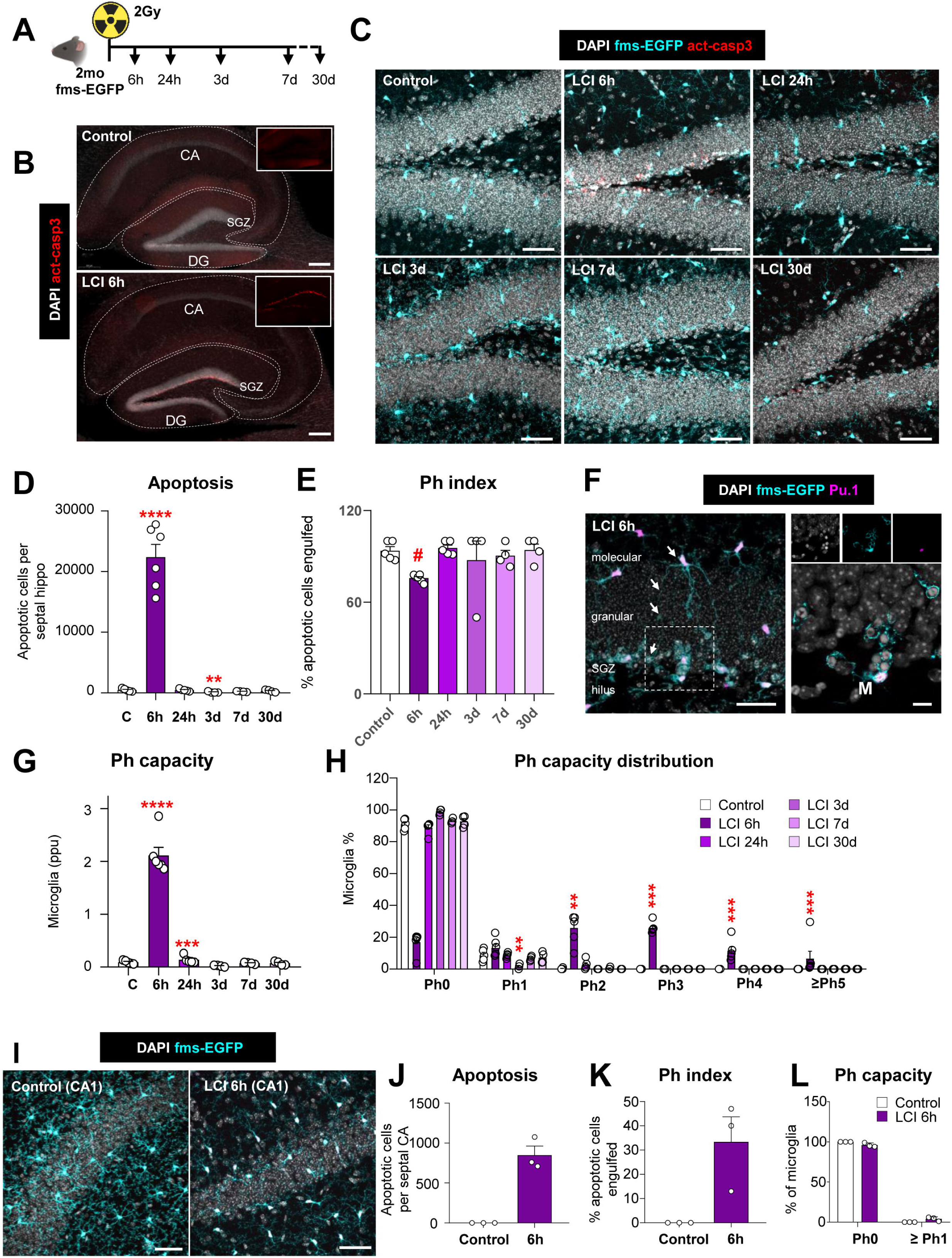
Low Cranial Irradiation (LCI) drives superphagocytic microglia. **(A)** Experimental design of superphagocytosis by LCI (2Gy). **(B)** Representative image of the hippocampus showing DAPI (white) and activated caspase 3 (act-casp3, red). The areas drawn with a dotted line differentiate the Cornus Ammonis (CA) and the dentate gyrus (DG), containing the subgranular zone (SGZ). **(C)** Representative confocal images of the LCI time course (6h-30d) showing nuclei with DAPI (white), apoptotic cells with activated caspase 3 (red, empty arrowheads), and microglia with fms-EGFP (cyan) staining. **(D)** Number of apoptotic cells in the DG per septal hippocampus. **(E)** Ph index (% of apoptotic cells phagocytosed by microglia) in the DG. **(F)** Representative images of superphagocytic microglia 6h after LCI showing the nuclear morphology with DAPI (white), microglial nuclei with Pu.1 (magenta), and microglia morphology with fms-EGFP (cyan) staining. White arrows indicate a microglial process that extends through the granular layer to perform phagocytosis and the cell in the dotted square is shown at higher magnification on the right panel. **(G)** Weighted Ph capacity of microglia (in parts per unit, ppu). **(H)** Ph capacity histogram (% of microglia containing phagocytic pouches) through the LCI time course. **(I)** Representative confocal images of CA in control and LCI 6h showing nuclear morphology with DAPI (white), and microglia with fms-EGFP (cyan) staining. **(J)** Number of apoptotic cells in the CA per septal hippocampus **(K)** Ph index (% of apoptotic cells phagocytosed by microglia) in CA. **(L)** Ph capacity histogram (% of microglia containing phagocytic pouches) in CA. Bars represent mean ± SEM of n=3-6 mice. Data was analyzed depending on the number of variables to be compared using t-test or one-way ANOVA followed by Holm-Sidak post hoc tests when appropriate. Some data [D, G, H] were Log transformed to comply with homoscedasticity. When homoscedasticity was not accomplished Mann Whitney test [J,K,L] or Kruskal-Wallis test [E,H] was performed.]. Asterisks represent significance between control and LCI (6h, 24h, 3d, 7d, 30d). ** represents p<0.01, *** represents p<0.001. # [E]=0.051. Scale bars= 200μm [B], 50μm [C], 20μm [F], 10μm (insert in [F]), 50μm [I]; z=1μm [B], z=21μm, z=17.5μm, z=25.2μm, z=13.3μm, z=16.1μm, z=14.7μm [C, left to right, top and bottom], z=13.3μm [F], z=4.2μm (insert in [F]), z=12.86μm, z=21μm [I, left to right].

We validated the use of LCI to assess post-phagocytic changes by determining microglial DNA damage and monocyte infiltration indicative of blood-brain barrier dysfunction. We used histone γH2AX to label double-strand breaks^28^ and found γH2AX-labeled SGZ nuclei but no microglial nuclei, identified with Pu.1^+^. Nonetheless, γH2AX puncta were found in microglial nuclei, although they occupied 85.8 ± 1.6% less nuclear area than in the rest of the SGZ nuclei (**Supp. Fig. 1A-E**). Infiltration of monocytes (fms-EGFP^+^, P2Y12R^-^) was undetectable across the time course (**Supp. Fig. 1F, G**), although we detected a small percentage of cells expressing *MS4a7*^29^ (Membrane Spanning 4-Domains A7) by RNAScope, which could be either infiltrating monocytes or border associated-macrophages (**Supp. Fig. 1H, I**), suggesting that LCI did not cause major microglial DNA breaks nor blood brain barrier damage, possibly because microglial DNA is densely packed^30^. Importantly, apoptosis and microglial phagocytosis were barely detected in the other main region of the hippocampus, the Cornu Ammonis (CA) (**Fig. 1J-L**), allowing us to use the DG *vs* CA comparison as an internal control to discriminate the effects of irradiation (occurring in both the DG and the CA) and the effects of phagocytosis (occurring only in the DG).

### Early post-phagocytic microglia display increased expression of galectin 3 and CD68

We then analyzed post-phagocytosis transcriptional changes by performing scRNA-Seq 24h and 7d after LCI (**Fig. 2**). The analysis pipeline included removal of reactive cells with artifact signature due to the processing^25^ and removal of macrophage and non-myeloid clusters prior to reembedding (**Sup. Fig. 2**). We identified 6 canonical microglia clusters: homeostatic (HM1, HM2), disease-associated (DAM), lysosomal (LM), interferon-response (IRM), and proliferative (PM). We found no changes in size either in HM1 or DAM, and a small size reduction in HM2 (**Supp. Fig. 2G**), possibly resulting from global effects of LCI, despite the low dose of irradiation used. However, the LM, the IRM, and the PM clusters increased 24h or 7d after LCI and could therefore be related to post-phagocytic microglia (**Fig. 2B, Supp. Fig. G**)

**Figure 2.**
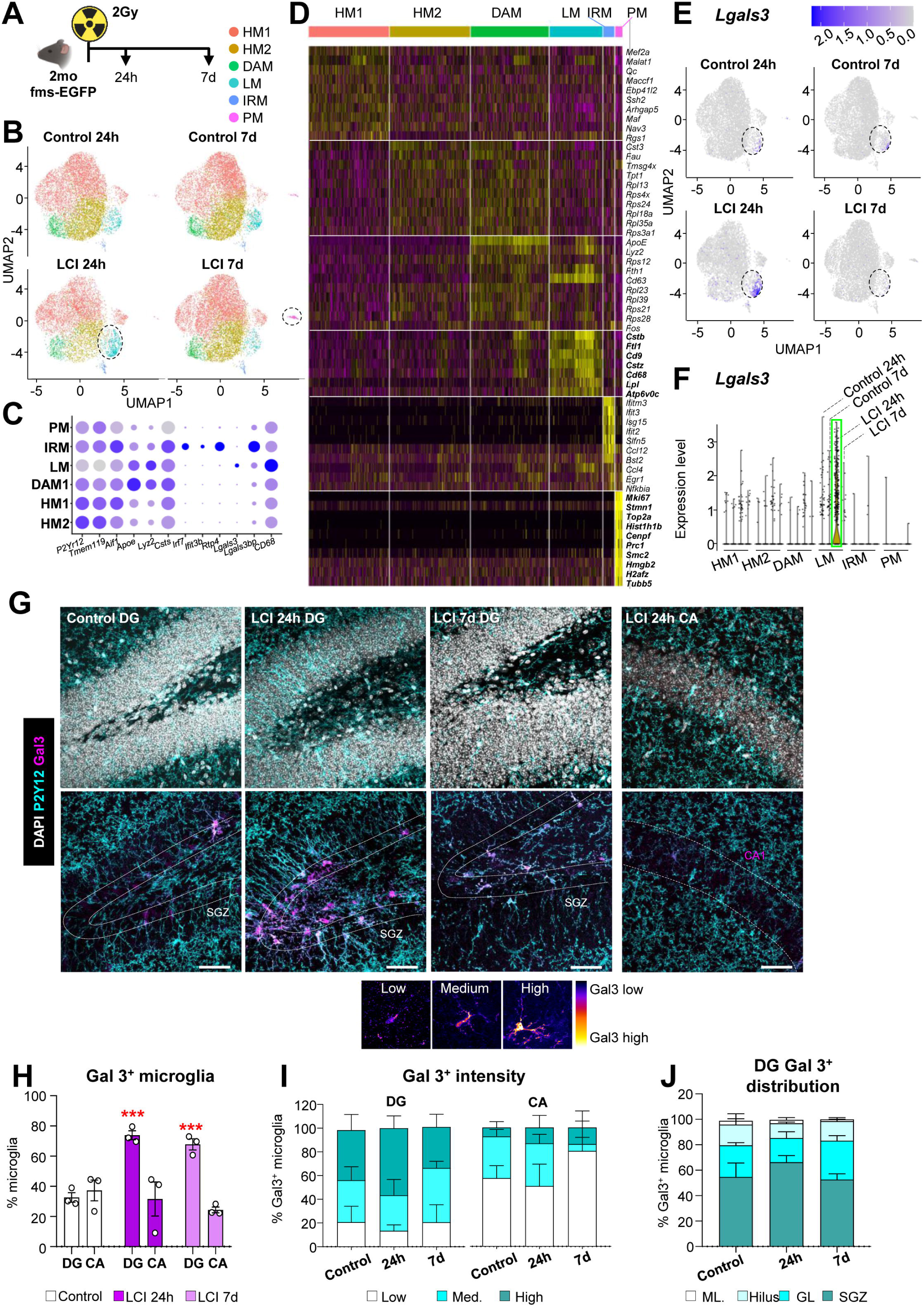
Early post-phagocytic microglia show increased expression of *Lgals3* and Gal3. **(A)** Timeline of the scRNA-seq experiment after LCI **(B)** UMAP visualization of microglial subpopulations identified through single-cell analysis: HM1 (homeostatic 1), HM2 (homeostatic 2), DAM (disease-associated microglia), LM (lysosomal microglia), IRM (interferon responsive microglia) and PM (proliferative microglia). Dotted black circles highlight increased LM cell numbers at 24 hours post-LCI and an increased PM cell numbers at 7 days post-LCI. **(C)** Dot plot illustrating top 5 enriched gene expression among the different microglia clusters. **(D)** Heatmap of top 10 differentially expressed genes for each microglia cluster. **(E)** UMAP showing the fold-change in *Lgals3* expression following LCI, with the LM subpopulation highlighted by a dotted black circle. **(F)** Violin plots depicting *Lgals3* expression across microglial clusters after LCI. **(G)** Representative confocal images of Gal3 immunofluorescence in the DG and CA after LCI, showing DAPI (white), Gal3 (magenta), and P2Y12 (cyan) staining. The SGZ is labelled with white dotted lines. The bottom image represents the classification criteria to define cells according to Gal3 intensity using a LUT (lookup table) from Image J. **(H)** Percentage of Gal3^+^ microglia in DG and CA after LCI. **(I)** Intensity of Gal3^+^ microglia in the DG and CA- **(J)** Topological distribution of Gal3^+^ microglia in the DG. Bars show mean ± SEM of n=3-4 mice. Data was analyzed by one-way ANOVA [H,I,J] followed by Holm-Sidak post hoc tests when appropriate. Asterisks represent significance between control and LCI (24h, and 7d). *** represents p<0.001. Scale bars=50μm. z=12.6, z=28.7, z=18.98, z=17.5 [F, left to right].

The LM cluster increased in size 24h after LCI (from 3.3 to 7.7% of total cells) and returned to basal levels by 7d (**Fig. 2B**, **Supp. Fig. 2G**). Among the top 10 genes expressed in the LM cluster were lysosomal genes such as *Cstb* and *Cstz* (cystatins B and Z), the lysosomal/endosomal-associated membrane glycoprotein *Cd68* (CD68 antigen), and enzymes such as *Ftl1* (ferritin light chain), *Cd9* (tetraspanin 29), *Lpl* (lipoprotein lipase) and *Atp6v0c* (ATPase H+ Transporting V0 Subunit C) (**Fig. 2C, D**). The LM cluster was characterized by specific expression of *Lgals3*, which encodes for galectin 3 (Gal3), a pleiotropic carbohydrate-binding protein involved in microglial responses to neurodegeneration^31^; and by increased expression of *Cd68* (**Fig. 2E, D**). To confirm *Lgals3* expression *in situ* we analyzed the expression of Gal3 by immunofluorescence. Gal3 expressing cells and intensity of expression increased specifically in post-phagocytic microglia in the DG 24 and 7d post-phagocytosis, but not in CA (**Fig. 2G, H, I**). Most of the DG Gal3^+^ microglia were located in the SGZ, supporting their relation to phagocytosis (**Fig. 2J**). To further confirm the relation of Gal3 with phagocytosis, we assessed its expression in microglia during early developmental stages in the hippocampus, from postnatal (P) day 2 to 28. In this period, we found a strong correlation (R^2^=0.752, p<0.001) between the % of Gal3^+^ expressing microglia and the engagement of the population in phagocytosis (**Fig. 3A, B**). As after LCI, during postnatal development most DG Gal3^+^ microglia were located in the SGZ neurogenic niche (**Fig. 3C**), where newborn cells undergo apoptosis^11^. Similarly, we analyzed the expression of CD68 by immunofluorescence and found increased expression in DG but not in CA microglia, confirming that the LM cluster was related to phagocytosis (**Fig. 3D-F**). However, unlike *Lgals3*, *Cd68* was not specific to the LM cluster but was expressed in many microglia regardless of their involvement in phagocytosis (**Fig. 2C, 3D; Supp. Fig 3**). We further studied changes in lysosomes using transmission electron microscopy and immuno-gold to identify microglia (**Fig. 3G-I**). In the scant microglial cytoplasm we found a large accumulation of phagosomes at 6h (**Fig. 3G**), and a non-significant trend towards an increased number of lysosomes containing large lipid droplets between 24h and 7d after LCI (ANOVA p value=0.101) (**Fig. 3H**). As Gal3 is involved in lysosomal repair and replacement^33^, the above results suggest that Gal3 and CD68 may be involved in a coordinated response to recover damaged or worn-out lysosomes, maintaining their availability in post-phagocytic microglia.

**Figure 3.**
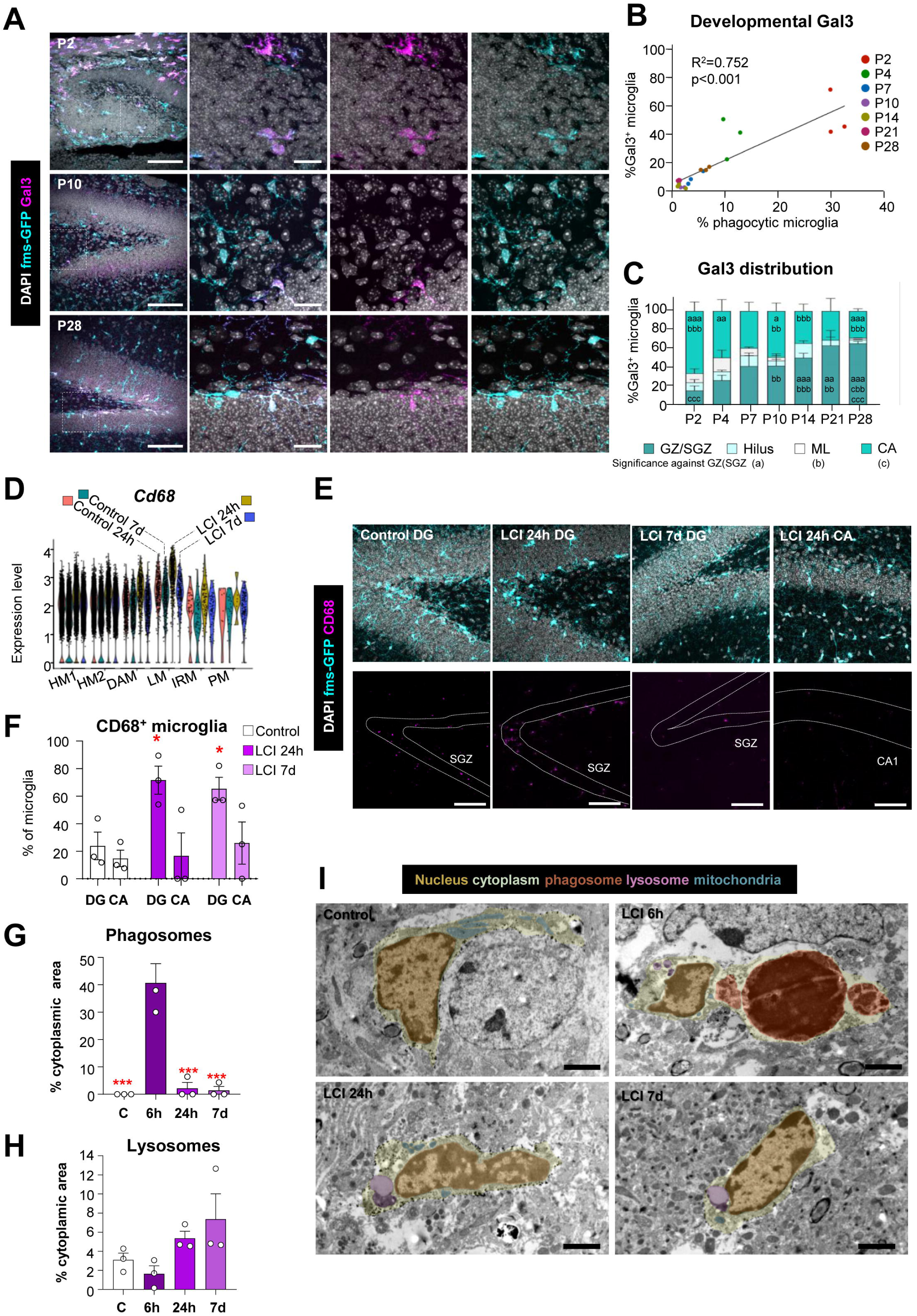
Early post-phagocytic microglia show increased expression of CD68 and increased lysosomes. **(A)** Representative confocal images of microglia expressing galectin3 in different developmental stages (postnatal days P2-P28), showing DAPI (white), Gal3 (magenta), and fms-EGFP (cyan) staining. Representative details are shown within dotted white squares. **(B)** Correlation of the percentage of Gal3^+^ microglia and the percentage of phagocytic microglia. **(C)** Topological distribution of Gal3^+^ microglia in the DG across development. **(D)** Violin plots showing *Cd68* expression across microglial clusters after LCI. **(E)** Representative confocal images of CD68 immunofluorescence in the DG and CA after LCI, showing DAPI (white), CD68 (magenta), and P2Y12 (cyan) staining. The SGZ is labelled with white dotted lines. **(F)** Percentage of CD68^+^ microglia in DG and CA after LCI. **(G)** Cytoplasmic microglial area occupied by phagosomes after LCI. **(H)** Cytoplasmic microglial area occupied by lysosomes after LCI. **(I)** Representative TEM images of the DG in control and after LCI Images show microglial cytoplasm (green) and microglial intracellular organelles, such as the nucleus (brown), phagosomes (orange), lysosomes (purple) and mitochondria (blue). Phagosomes were mostly found at 6h. Bars represent mean ± SEM of n=3-4 mice [C,F,G,H]. Data was analyzed by one-way ANOVA followed by Holm-Sidak post hoc tests when appropriate [C,F,G,H]. Asterisks represent significance between control and LCI (24h, and 7d) [F] and between all the experimental conditions [G,H]. * represents p<0.05 and *** represents p<0.001. Scale bars= 50μm [A], 50μm [E], 2μm [I]; z stack de [A], z=18.2, z=8.4, z=23.1, z=17.5 [E, left to right].

Another potential cluster that could harbor the post-phagocytic microglia population was the IRM cluster, recently involved in phagocytosis^5^, whose relative size also increased 24h after LCI (from 0.54 to 1.32%; **Supp. Fig. 2G**). As a marker of IRM we analyzed *Ifit3* (Interferon Induced Protein With Tetratricopeptide Repeats 3)^5^ using RNAScope. However, we did not observe P2Y12⁺ microglial cells displaying robust *Ifit3* labeling. Nonetheless, we quantified the percentage of microglia showing a sparse, punctuated expression pattern, which remained unchanged in both the DG and CA regions at 24 hours and 7d post-LCI (**Supp. Fig. 4A, B**). We also analyzed by immunofluorescence another marker of the IRM signature, IFITM3 (Interferon Induced Transmembrane Protein 3)^5^, but it was barely detectable in the hippocampus at any time point after LCI, and it was only found in blood vessels (**Supp. Fig. 4C**), possibly in endothelial cells^34^. These results suggest that the IRM cluster did not contain post-phagocytic microglia.

To further characterize the post-phagocytic microglia after superphagocytosis, we analyzed the expression of the homeostatic marker P2Y12 and found it clustered in phagocytic pouches 6h after LCI (**Fig. 4A-C, Supp. Fig. 4D**), possibly because of its recruitment during the recognition of apoptotic cells. Finally, we used Sholl analysis to identify morphological changes in DG microglia but did not detect major morphological changes at 24h and 7d after LCI (**Fig. 4D-F**). However, microglia engaged in phagocytosis (i.e., with phagocytic pouches) at 6h after LCI did show reduced morphological complexity characterized by reduced number of intersections, suggestive of reduced number of processes (**Fig. 4G-I**). Thus, microglia engaged in phagocytosis displays phagocytic pouches and reduced morphological complexity, with no major changes observed at later time points

**Figure 4.**
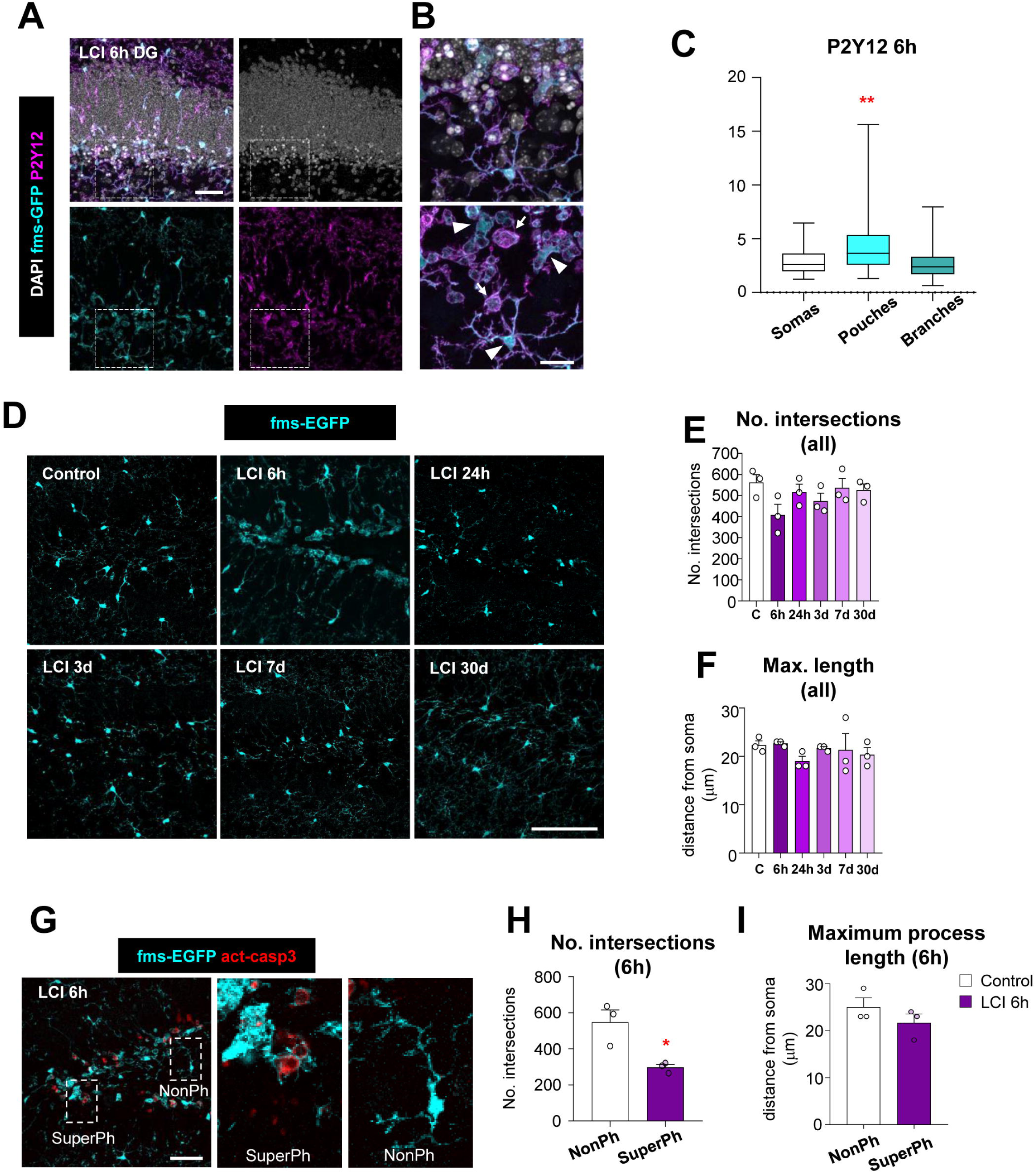
Early post-phagocytic microglia show P2Y12 accumulation in phagocytic pouches and a simplified morphology. **(A)** Representative images of phagocytic pouches in the DG 6h after LCI showing DAPI (white), P2Y12 (magenta), and fms-EGFP (cyan) staining. **(B)** Details of P2Y12 expression in the microglial soma (white arrowhead) and phagocytic pouches (white arrows) showing DAPI (white), P2Y12 (magenta), and fms-EGFP (cyan) staining. **(C)** Box-plot representing the expression level of P2Y12 in the soma and phagocytic pouches of microglia 6h afterLCI. **(D)** Representative confocal images of microglial morphology showing fms-EGFP (cyan) after LCI. **(E, F)** Sholl analysis of microglial morphology after LCI showing the total number of intersections, representative of the total number of processes **(E)**, and the maximum distance of the processes **(F)**. **(G)** Representative detail of superphagocytic and non-phagocytic microglia at LCI 6h showing activated caspase 3 (red) and fms-EGFP (cyan) staining. **(H, I)** Sholl analysis of microglial morphology 6h after LCI comparing superphagocytic and non-phagocytic cells showing the total number of intersections, representative of the total number of processes **(H)**, and the maximum distance of the processes **(I)**. Bars represent the mean ± SEM of 3-4 independent experiments. Data was analyzed by one-way ANOVA followed by Holm-Sidak post hoc tests when appropriate [E,F], by Kruskal-Wallis analysis when data did not meet normality and homoscedasticity [C] and by Student’s t-test [H,I]. Asterisks represent significance between groups. * represents p<0.05 and ** represents p<0.01. Scale bars=50μm [A,D,G], 20μm [B]; z=18.9 [A]; z=18.9 [B]; z=21μm, z=15.4μm, z=18,9; z=8.4μm, z=14μm, z=12.6μm [D, left to right, top and bottom]; z=11.9 [G].

### Late post-phagocytic microglia undergo abortive proliferation

We then explored the changes in the PM cluster 7d after LCI, and confirmed the increased expression of S and G2M phase genes (**Fig. 5A, B**). To determine whether proliferation was specific to post-phagocytic microglia or a response to LCI, we used Ki67 (**Fig. 5C, D**), a product of *Mki67* (**Fig. 2D**), whose levels increase during S phase and reach a maximum during G2/M^35^. Proliferative cells in the DG (presumably neuroprogenitors) were wiped off at 6h, and recovered gradually between 3 and 30d (**Fig. 5E**). Proliferation of microglia was induced between 7d and 30d after LCI specifically in the DG (**Fig. 5F,G**), suggesting that it was not related to LCI but to post-phagocytosis events. We speculate that this proliferation was possibly triggered to compensate for the decrease in the number of microglia found at 24h after LCI, as it did not lead to increased microglia number over the time course (**Fig. 5H**). To further confirm induction of proliferation in post-phagocytic microglia we injected mice with the thymidine analog 5-bromo-2’-deoxyuridine (BrdU) 7d after LCI and analyzed its expression 24h and 48h later (**Fig. 5I-K**). We found that whereas total BrdU^+^ cells remained constant in the 24h-48h window, BrdU^+^ microglia were significantly lost (**Fig. 5L, M**), suggesting an abortive microglial proliferation program triggered by phagocytosis.

**Figure 5.**
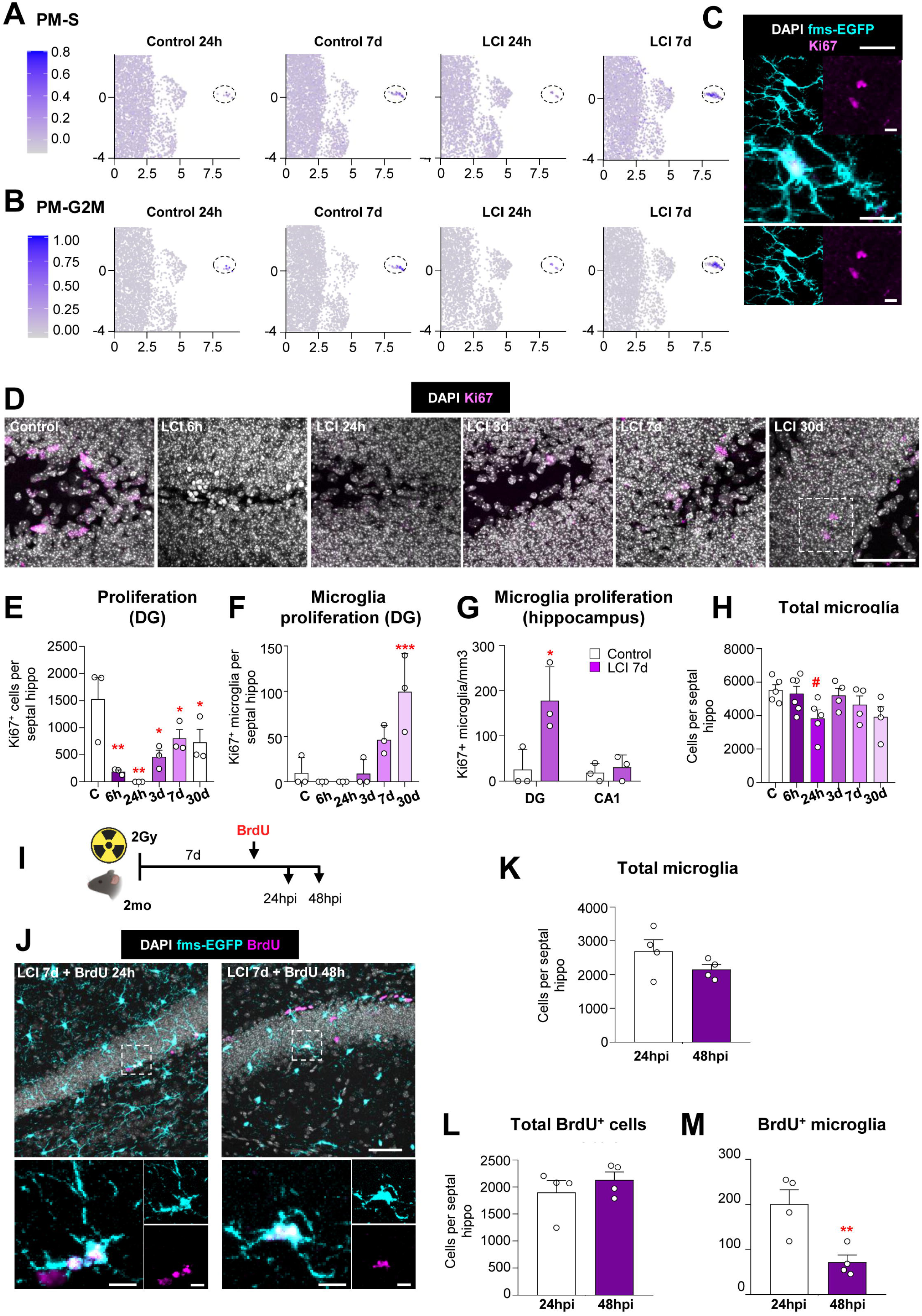
Late post-phagocytic microglia display abortive proliferation. **(A)** UMAP illustrating the enrichment score of S-phase gene in the PM cluster (highlighted by a dotted black circle) after LCI. **(B)** UMAP illustrating the enrichment score of G2/M-phase gene in the PM cluster (highlighted by a dotted black circle) after LCI. **(C)** Detail of proliferative microglia showing DAPI (white), Ki67 (magenta), and fms-EGFP (cyan) staining 30d after LCI. **(D)** Representative confocal images of proliferation showing DAPI (white), Ki67 (magenta), and GFP (cyan) staining after the LCI time course. The pair of dividing cells at 30d (dotted white box) is shown in Fig. 5C. **(E)** Number of Ki67^+^ cells in the DG per septal hippocampus. **(F)** Number of Ki67^+^ microglia in the DG per septal hippocampus. **(G)** Density of Ki67^+^ microglia in the DG and CA. **(H)** Number of microglia in the DG per septal hippocampus. **(I)** Experimental design of BrdU assay after LCI. BrdU was injected 7d after LCI and microglial proliferation was assessed 24 and 48h post-BrdU injection (hpi). **(J)** Representative confocal images of newly born microglia at 24 and 48hpi showing DAPI (white), BrdU (magenta) and fms-EGFP (cyan) staining. The dotted square indicated by an arrow shows details of BrdU^+^ microglia. **(K)** Number of microglia in the DG per septal hippocampus. **(L)** Number of BrdU^+^ cells in the DG per septal hippocampus. **(M)** Number of BrdU^+^ microglia in the DG per septal hippocampus. Bars show mean ± SEM of n=3-4 mice. Data was analyzed by one-way ANOVA [E,F,H] followed by Holm-Sidak post hoc tests when appropriate, and by Student’s t-test [G,K,L,M]. Asterisks represent significance between control and LCI (6h, 24h, and 7d) [E,F,H], between control and LCI 7d [G] and between 24hpi and 48hpi [M]. * represents p<0.05, ** represents p<0.01, *** represents p<0.001; # represents p=0.051. Scale bars= 10μm [C], 50μm [D,J], 10μm (inserts in [J]). z=16.1μm [C]; z=8.4μm, z=11.2μm, z=16.1μm, z=14.7μm, z=16.1μm, z=16.1μm [D, left to right]; z=13.3μm, z=27.3μm [J, left to right].

### Phagocytosis triggers microglial death, oxidative stress, and metabolic rewiring

We hypothesized that proliferation in post-phagocytic microglia was the result of phagocytosis-induced microglial death. We observed examples of phagocytic and post-phagocytic microglia undergoing apoptosis after LCI (**Fig. 6A, B**), but the phenomenon was too rare to be quantified, possibly because apoptotic microglia were engulfed by nearby microglia. To understand the mechanisms underlying phagocytosis-induced microglial death, we used *an in vitro* model of phagocytosis in which microglia were fed with apoptotic SH-SY5Y neurons (pretreated with staurosporine) (**Fig. 6C**). We detected many transcriptional changes in post-phagocytic microglia (24h) (original gene array data published in ^10^), which included genes in Gene Ontologies Apoptosis and Proliferation (including genes shared by the two GO terms) (**Fig. 6D**). In this *in vitro* model, we detected a consistent increase in microglial apoptosis throughout the 24h time course (**Fig. 6E**), and a significant increase in the production of intracellular reactive oxygen species (ROS) (**Fig. 6F,G**). Phagocytosis-induced microglial apoptosis was prevented by pretreatment with the antioxidant N-acetyl-cysteine (NAC) at 0.1 and 1mM, but was exacerbated with 10mM NAC (**Fig. 6H**), possibly related to toxic dose-dependent effects^36^. NAC did not affect phagocytosis efficiency (**Supp. Fig. 5A**), confirming that its protective effects at low concentrations were due to a reduction of the oxidative stress induced by phagocytosis.

**Figure 6.**
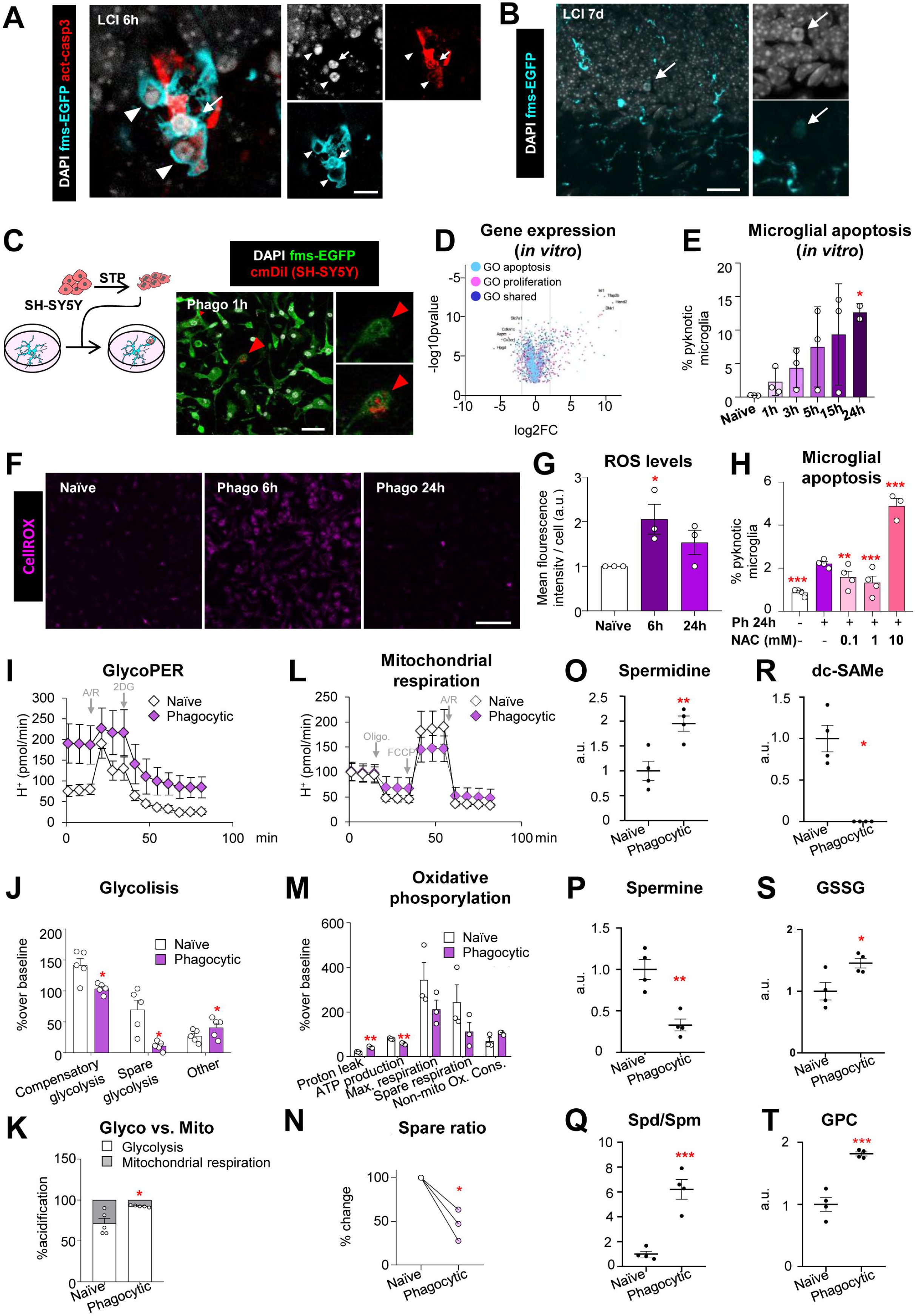
Phagocytosis triggers apoptotic cell death, oxidative stress, and metabolic rewiring. **(A)** Representative confocal image of phagocytic microglia undergoing apoptosis 6h after LCI showing DAPI (white), activated caspase 3 (red) and fms-EGFP (cyan) staining. The arrow points to the pyknotic, activated caspase3+ nucleus of microglia, and the arrowheads to the phagocytic pouches. Note that phagocytic pouches, unlike the microglial nucleus, do not contain fms-EGFP staining. **(B)** Representative confocal image of phagocytic microglia undergoing apoptosis 6h after LCI, showing DAPI (white) and fms-EGFP (cyan) staining. **(C)** Experimental design of *in vitro* phagocytosis assay using Vampire-SH-SY5Y cells pretreated with staurosporine (3μM) and representative confocal images of microglial phagocytosis at 1h showing DAPI (white), Vampire-SH-SY5Y (red) and fms-EGFP (green) staining. The detail shows SH-SY5Y content within microglia. The images were originally printed in ^10^ and are reprinted with permission. **(D)** Volcano plot showing genes related to the GO terms apoptosis, proliferation, or both in a gene microarray comparing naïve vs phagocytic (24h) microglia from the in vitro model of phagocytosis (data originally published in ^10^). **(E)** Percentage of pyknotic microglia after phagocytosis *in vitro*. **(F)** Representative confocal images of cultured naïve and phagocytic microglia (6h, 24h) expressing ROS detected by CellROX. **(G)** ROS levels indicated in arbitrary units (a.u.). **(H)** Percentage of pyknotic microglia after phagocytosis *in vitro* (24h) and pre-treatment with the antioxidant N-Acetylcysteine (NAC). **(I)** Representative experiment showing the proton extrusion rate (PER) in pmol/min caused by glycolysis in naïve and phagocytic microglia (24h). **(J)** Analysis of glycolysis (PER normalized to baseline) in naïve and phagocytic microglia. **(K)** Percentage of PER originated by glycolysis and oxidative phosphorylation in naïve and phagocytic microglia. **(L)** Representative experiment showing the oxygen consumption rate (OCR) in pmol/min of naïve and phagocytic microglia (24h) **(M)** Analysis of oxidative phosphorylation (OCR normalized to baseline) in naïve and phagocytic microglia. **(N)** Normalized spare capacity of mitochondrial respiration in naïve and phagocytic microglia. **(O)** Relative production of spermidine in arbitrary units (a.u.) in naïve and phagocytic microglia. **(P)** Relative production of spermine in arbitrary units (a.u.) in naïve and phagocytic microglia. **(Q)** Sermidine/spermine ratio in naïve and phagocytic microglia. **(R)** Relative production of dc-SAMe in arbitrary units (a.u.) in naïve and phagocytic microglia. **(S)** Relative production of GSSG in arbitrary units (a.u.) in naïve and phagocytic microglia. **(R)** Relative production of GPC in arbitrary units (a.u.) in naïve and phagocytic microglia. Bars show mean±SEM of 3-6 independent experiments [E,G,H,J,M,K]. Dots represent individual measurements [N-T]. Icons (diamonds) represent the mean of the different conditions of a representative experiment, while error bars show the SD [I,L]. Data was analyzed by one-way ANOVA [E,G,H] followed by Holm-Sidak post hoc tests [E,G] and Dunnet’s test [H] when appropriate. Data was analyzed by Student’s t-test [J,K,M,O,P,Q,S,T]. Some data [J,M] was log-transformed to comply with homoscedasticity When neither normality nor homoscedasticity were met, the Mann-Whitney test performed [N,R]. * indicates p <0.05. ** indicates p <0.01, *** indicates p <0.001. Scale bars= 30μm [A,C], 30μm [B], 30μm [F]. z=6.3μm [A]; z=0.7μm [C]; z=0.7μm [F];

To understand the mechanisms underlying the phagocytosis-induced oxidative stress, we further analyzed the transcriptional changes induced by phagocytosis in culture^10^ using WGCNA (weighted correlation network analysis (**Supp. Fig. 5B, C**).). We identified four expression modules at 3 and 24h after phagocytosis (up, down, transient up, transient down), in agreement with our previous analysis^10^. The Up showed strongly significant GOs related to neurogenesis and metabolism, while the Up-Down module (transient upregulated, then downregulated) displayed a large number of significantly changed GO terms associated with cell cycle (**Supp. Fig. 5B, C**). We then looked at individual genes involved in key catabolic pathways in this set of genes and found a consistent reduction in multiple genes related to oxidative phosphorylation (pyruvate mitochondrial transport, acetylCoA synthesis, Krebs cycle, and electron transport chain), but not a clear transcriptional effect on genes related to glycolysis, which included both upregulated and downregulated genes (**Supp. Fig. 5D**).

Next, we directly assessed glycolysis and oxidative phosphorylation using Seahorse analysis of the proton flux under different metabolic inhibitors, in microglial cultures grown at a physiological glucose concentration (2.5mM) fed with apoptotic SH-SY5Y neurons. Post-phagocytic microglia at 24h showed a reduction in those two major catabolic pathways that resulted in a switch towards glycolysis (**Fig. 6I-M**) that is reminiscent, but not identical, to the Warburg effect found during phagocytosis-induced trained immunity in other macrophages^37^. Nonetheless, post-phagocytic microglia showed decreased compensatory and spared glycolysis (**Fig. 6I, J**), and decreased mitochondrial ATP production (**Fig. 6L, M**) that led to reduced mitochondrial spare respiratory capacity, defined as the difference between basal ATP production and its maximal activity (**Fig. 6N**). Accordingly, phagocytic microglia showed reduced mitochondrial complexity compared to naïve cells in culture, using both BV2 cells and primary cultures. In BV2 microglia transfected with MitoGFP, we analyzed by confocal microscopy the 3D mitochondrial network and observed that post-phagocytic microglia showed reduced both individual mitochondria and mitochondrial networks, with a trend towards reduced length, branches, and junctions, but no significant changes in other morphological parameters compared to naïve cells (**Supp. Fig. 6A-C**). We confirmed these results in primary microglia analyzed by electron microscopy, and observed reduced number of mitochondrial objects that occupied less cytoplasmic area but no changes in microglial morphology in post-phagocytic microglia compared to naïve cells (**Supp. Fig 6D-G**). Altogether, these morphological alterations could be responsible for the decreased oxidative phosphorylation in post-phagocytic microglia.

The Seahorse analysis detected that other metabolic pathways could be increased in post-phagocytic microglia at 24h (**Fig. 6J**). We tested this possibility using targeted metabolomics with mass spectrometry for metabolites of the methionine and folate cycles (**Fig. 6O-S; Supp. Fig. 7**), some of which are related to oxidative stress. Cultured post-phagocytic microglia at 24h displayed changes in polyamine synthesis, with increased spermidine levels and reduced spermine levels that led to a higher spermidine/spermine ratio (**Fig. 6O-Q**). These polyamines are involved in the recovery from oxidative stress, e.g., increased spermidine mitigates oxidative stress by reducing ROS levels^38^, cell proliferation, and differentiation^39^. Post-phagocytic microglia also showed reduced decarboxylated S-adenosylmethionine (dc-SAMe) (**Fig. 6R**), a spermidine precursor, suggesting dc-SAMe exhaustion towards spermidine production; and increased glutathione disulfide (GSSG)(**Fig. 6S**), the oxidized form of glutathione (GSH; **Supp. Fig. 7A-C**), which is generated when glutathione acts as an antioxidant^40^. Additionally, phagocytosis also increased levels of the metabolite glycerylphosphorylcholine (GPC) (**Fig. 6T**), a precursor of phosphatidylcholine, a major component of cellular membranes in the body. The metabolite serine, which is involved in protein synthesis, in the folate cycle, and in glutathione synthesis, showed a trend towards increased production in post-phagocytic microglia (**Supp. Fig. 7D**), but metabolites of the Krebs cycle, such as succinate, malate and fumarate did not change significantly (**Supp. Fig. 7E**). Altogether, the metabolomic data suggested that after phagocytosis microglia synthetized metabolites involved in oxidative stress resolution, in cell proliferation, and in cell recovery, by the generation of proteins and lipidic membranes that may have been degraded during the phagocytic process-

### Phagocytosis efficiency is preserved in post-phagocytic microglia

The phagocytosis-induced oxidative stress and metabolic rewiring shown above suggested that microglial functionality may be compromised after phagocytosis. To test this hypothesis, we challenged microglia with two consecutive doses of LCI, 7 days apart (**Fig. 7A-E**). A shorter interval was not feasible because the proliferative cells in the neurogenic niche were wiped out at earlier time points after LCI (**Fig. 5E**). After the second challenge, we found ample evidences of microglial phagocytosis (**Fig. 7C**), whose efficiency was maintained, as determined as the Ph index (**Fig. 7D**). However, the second LCI challenge induced a substantially smaller amount of apoptosis (**Fig. 7E**), preventing the comparison of phagocytosis efficiency between the first and second LCI challenge.

**Figure 7.**
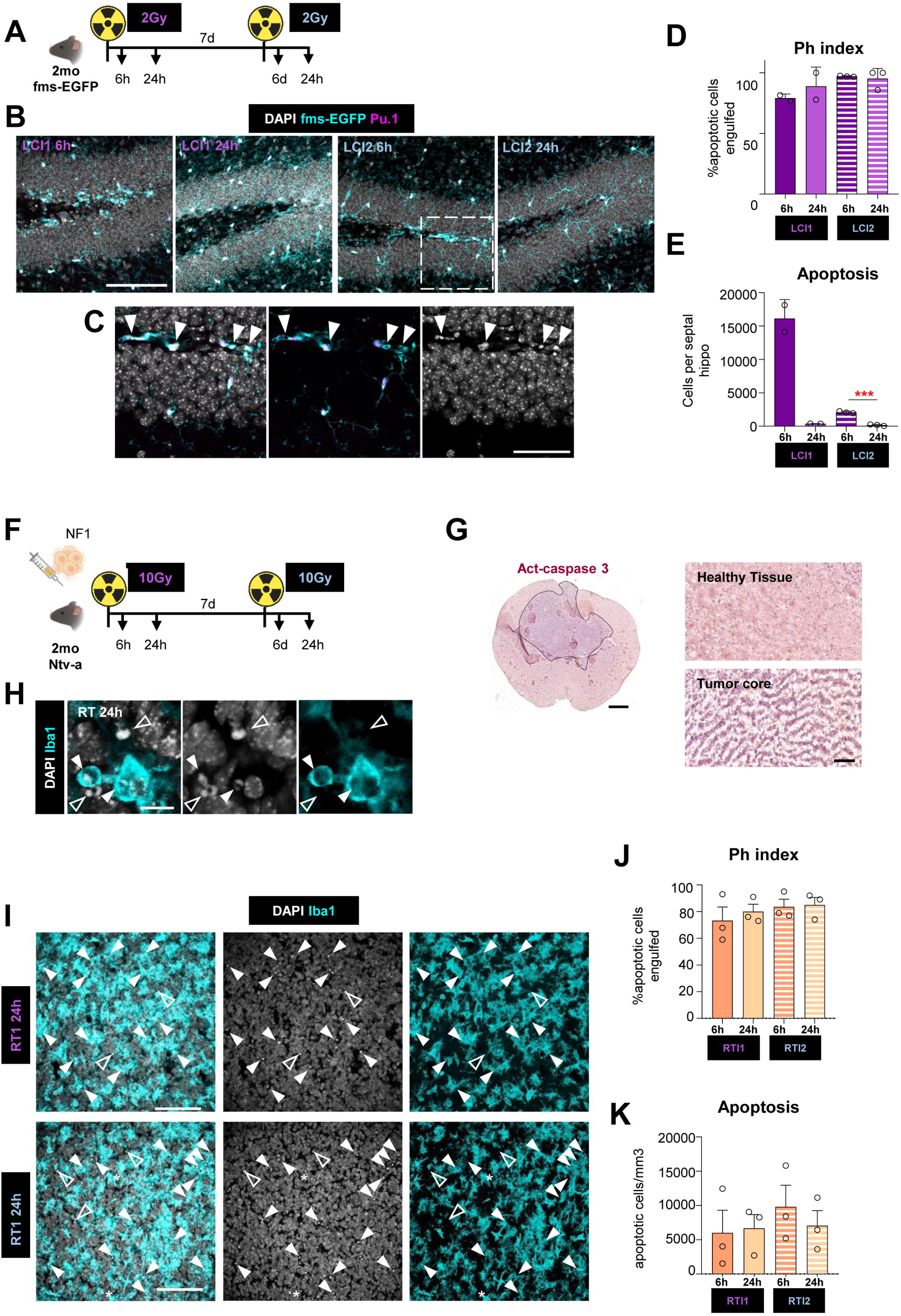
Microglial phagocytosis efficiency is preserved in the long-term after LCI and irradiated murine glioblastoma. **(A)** Experimental design to test the effect of consecutive LCI challenges (2Gy each) 7 days apart, analyzed at 6h and 24h in Nf1-silenced murine glioblastoma. **(B)** Representative confocal images of the different experimental conditions (LCI1 at 6h and 24h, and LCI2 at 6h and 24h) showing DAPI (white), Pu.1 (magenta) and fms-EGFP (cyan) staining. The cells in the white dotted box are shown in C. **(C)** Detail of Pu.1^+^ microglia engulfing pyknotic nuclei 6h after LCI2. The phagocytic pouches are indicated with white arrowheads. **(D)** Ph index (% of apoptotic cells phagocytosed by microglia) after LCI1 and LCI2. **(E)** Number of apoptotic cells in the DG per septal hippocampus after LCI1 and LCI2. **(F)** Experimental design of a mouse model (Ntv-a) bearing the Nf1-silenced glioblastoma to test the effect of consecutive radiotherapy (RT) challenges (10Gy each) 7 days apart, analyzed at 6h and 24. **(G)** Representative scanner image of a brain slice bearing Nf1-silenced glioblastoma immunostained for activated caspase 3, showing staining in the tumor core. **(H)** Representative confocal image of a phagocytic TAM (tumor-associated macrophage) exposed to RT at 24h showing DAPI (white) and Iba1 (cyan). White arrowheads show engulfed apoptotic cells and arrowheads with a white border show non-phagocytosed apoptotic cells. **(I)** Representative confocal images of the NF1 glioblastoma 24h after RT1 and RT2 showing DAPI (white) and Iba1 (cyan) staining. White arrowheads show engulfed apoptotic cells and arrowheads with a white border show non-phagocytosed apoptotic cells. **(J)** Ph index (% of apoptotic cells inside the tumor phagocytosed by TAMs) after RT1 and RT2 in the tumor core. **(K)** Density of apoptotic cells inside after RT1 and RT2 in the tumor core. Bars show the mean ± SEM of n=3 mice (n=2 for positive controls). Data was analyzed by Student’s t-test when comparing time-dependent differences (6h vs 24h) [C,D,L,M]. Data was analyzed by one-way ANOVA [D, E, J, K] followed by Holm-Sidak post hoc tests when appropriate. * indicates p <0.05 and *** indicates p <0.001. Scale bars= 100μm [B, I], 10μm [H]. z=16.1μm, z=18.2μm, z=17.5μm, z=13.3μm [B, left to right], z=0.7μm [C]; z=0.7μm [IH; z=16.1μm and z=14μm [J, left to right].

Thus, we tested the functional impact of phagocytosis on a model of glioblastoma, the most aggressive form of brain cancer, which is usually treated with sequential (hypofractionated) radiotherapy (RT). We used a model of the mesenchymal subtype-Nf1-silenced of glioblastoma using the RCAS-TVA system which uses avian leukosis virus (ALV)-based vectors to selectively infect cells containing the TVA receptor ^41^ (**Fig. 7F-G**). Phagocytosis by tumor-associated microglia and macrophages (TAMs) was identified through the presence of phagocytic pouches containing pyknotic nuclei. (**Fig. 7H**). Sequential doses of RT one week apart induced similar numbers of apoptotic tumor cells and a similar TAM phagocytosis efficiency (**Fig. 7I-K**), suggesting that the metabolic and transcriptional changes found in post-phagocytic microglia were adaptive and destined to sustain their phagocytosis efficiency in the long term.

### The regenerative properties of post-phagocytic microglia depends on galectin 3

Finally, we tested the impact of the phagocytosis-induced transcriptional adaptations on microglia and the surrounding neurogenic niche in Gal3 knock-out (KO) mice, 24h and 7d after LCI (**Fig. 8**). While Gal3 KO mice show deficits in immune activation^42^, we did not observe any effects on microglial survival and only a small reduction in phagocytosis efficiency 24h after LCI (**Fig. 8A-C**). In addition, Gal3 KO mice showed reduced number of apoptotic newborn cells through the time course, an effect that was more robust 24h after LCI (**Fig. 8D**). As apoptotic cells in the hippocampus are presumably newborn neurons undergoing apoptosis^11^, we then assessed whether neurogenesis was affected in Gal3KO mice. We found that Gal3KO mice failed to recover adult hippocampal neurogenesis 7d after LCI, determined by the number of young neuroblasts born after the LCI, labeled with doublecortin and showing an immature morphology (AB cells; **Fig. 8A, E, F**). While a significant effect of the genotype on mature neuroblasts born before LCI (CD and EF cells; **Supp. Fig. 8A**) was not detected, Gal3KO mice showed an overall reduction in total neuroblasts at all times tested (**Fig. 8A, G**), demonstrating the pro-neurogenic role of Galectin 3. In agreement with their reduced young neuroblast production, Gal3KO mice failed to recover proliferative cells in the neurogenic niche, labeled with Ki67, from 24h to 7d after LCI (**Fig. 8G, I, J; Supp. Fig. 8B**). In addition, Gal3KO had a significantly smaller recruitment of radial neuroprogenitors (rNPCs) engaged in proliferation at 7d after LCI (**Fig. 8I, K**), despite similar numbers of rNPCs across groups (**Fig. 8L**). Although a non-significant trend (p=0.055) towards reduced numbers of rNCPs in Gal3KO compared to WT mice suggested potential differences in basal neurogenesis that should be explored in the future, these results underscore the regenerative role of Gal3 expressed in post-phagocytic microglia by promoting rNPC proliferation and newborn neuron production.

**Figure 8.**
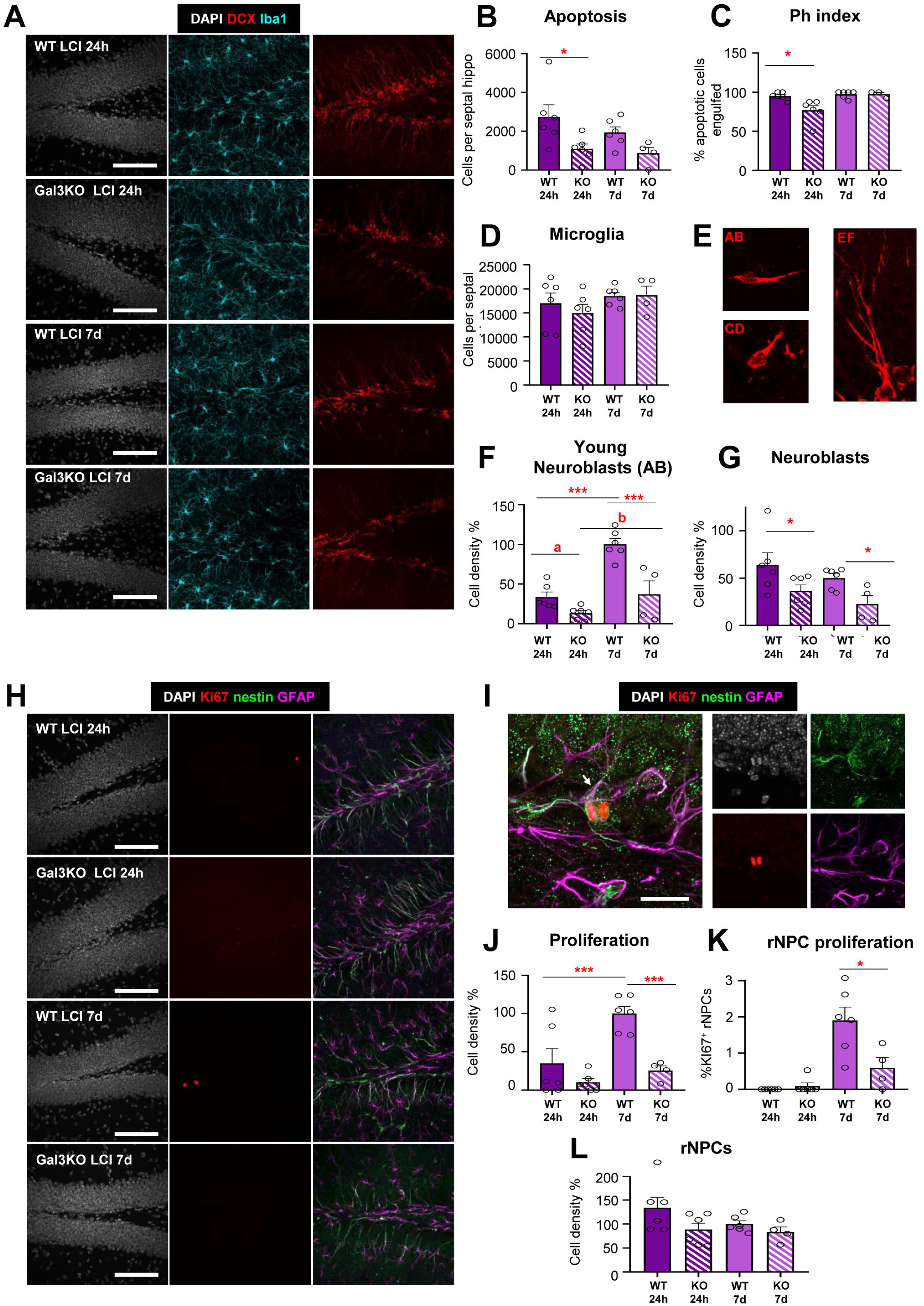
Galectin 3 deficient mice fail to recover the neurogenic niche after LCI. **(A)** Representative images of the hippocampus showing DAPI (white), microglia labeled with Iba1 (cyan) and neuroblasts labeled with doublecortin (DCX, red). **(B)** Density of microglia in the DG per septal hippocampus. **(C)** Density of apoptotic cells in the DG per septal hippocampus. **(D)** Ph index (% of apoptotic cells phagocytosed by microglia) in the DG. **(E)** Types of neuroblasts labeled with doublecortin (red) **(F)** Relative density of young (AB) neuroblasts in the DG per septal hippocampus. **(G)** Relative density of total neuroblasts in the DG per septal hippocampus. **(H)** Representative image of the hippocampus showing DAPI (white), the proliferation marker Ki67 (red), and the radial processes of the rNPCs labeled with both nestin (green) and GFAP (magenta). **(I)** High magnification confocal image of a proliferating rNPC in telophase, showing condensed chromosomes (DAPI, white) labeled with Ki67 (red), and an apical process labeled with nestin (green) and GFAP (magenta) indicated by an arrow. **(J)** Relative density of Ki67^+^ cells in the DG per septal hippocampus. **(K)** Percentage of Ki67^+^ rNPCs in the DG per septal hippocampus. **(L)** Relative density of rNPCs in the DG per septal hippocampus. Bars show the mean ± SEM of n=4-6 mice. Neurogenesis-related parameters (Ki67^+^ cells, rNPCs, and neuroblasts), which rely strongly on the age of the animal ^11^, were analyzed as relative densities compared to the WT group 7 days after LCI) in each of the two batches of animals analyzed (each comprising the four experimental groups). Data was analyzed by one-way ANOVA followed by Holm-Sidak post hoc tests when appropriate. * indicates p <0.05, ** indicates p<0.01, *** indicates p <0.001, a indicates p=0.054 and b indicates p00.06. Scale bars=100μm [A, H], 20μm [H]. z=18.2, 20.3, 23.1, 17.5 μm [A, top to bottom]; z=7μm [I]; z=16.8, 20.3, 16.8, 16.8 μm (G, top to bottom).

## DISCUSION

Here we set out to understand the time-dependent adaptations endured by microglia after phagocytosing apoptotic cells, and their functional consequences (**Figure 9**). We developed an *in vivo* model of superphagocytosis using LCI that synchronized DG microglia in engulfment by 6h, with cells displaying multiple pouches dotted with clusters of P2Y12R and reduced morphological complexity. By 24h, microglia had removed all apoptotic debris with an effective clearance time of 41 min per apoptotic cell. At this stage, post-phagocytic microglia were morphologically indistinguishable from non-phagocytic cells, but exhibited a lysosomal transcriptional signature characterized by increased expression of Gal3 and CD68, a trend towards increased number of lysosomes, and signs of apoptosis. We confirmed the phagocytosis-induced stress *in vitro*, showing that post-phagocytic microglia suffered oxidative stress, apoptosis, mitochondrial remodeling catabolic shutdown, and increased production of anti-oxidant polyamines. At later time points (7 days), microglia displayed a proliferative transcriptional signature but their proliferation was abortive and did not result in an increased cell number. These metabolic and transcriptional alterations were adaptive and allowed microglia to maintain their phagocytosis efficiency after LCI and in a model of irradiated glioblastoma. Finally, we used Gal3 deficient mice to show that the adaptations in post-phagocytic microglia were essential to recover the neurogenic niche after irradiation, underscoring the regenerative potential of microglial phagocytosis.

**Figure 9.**
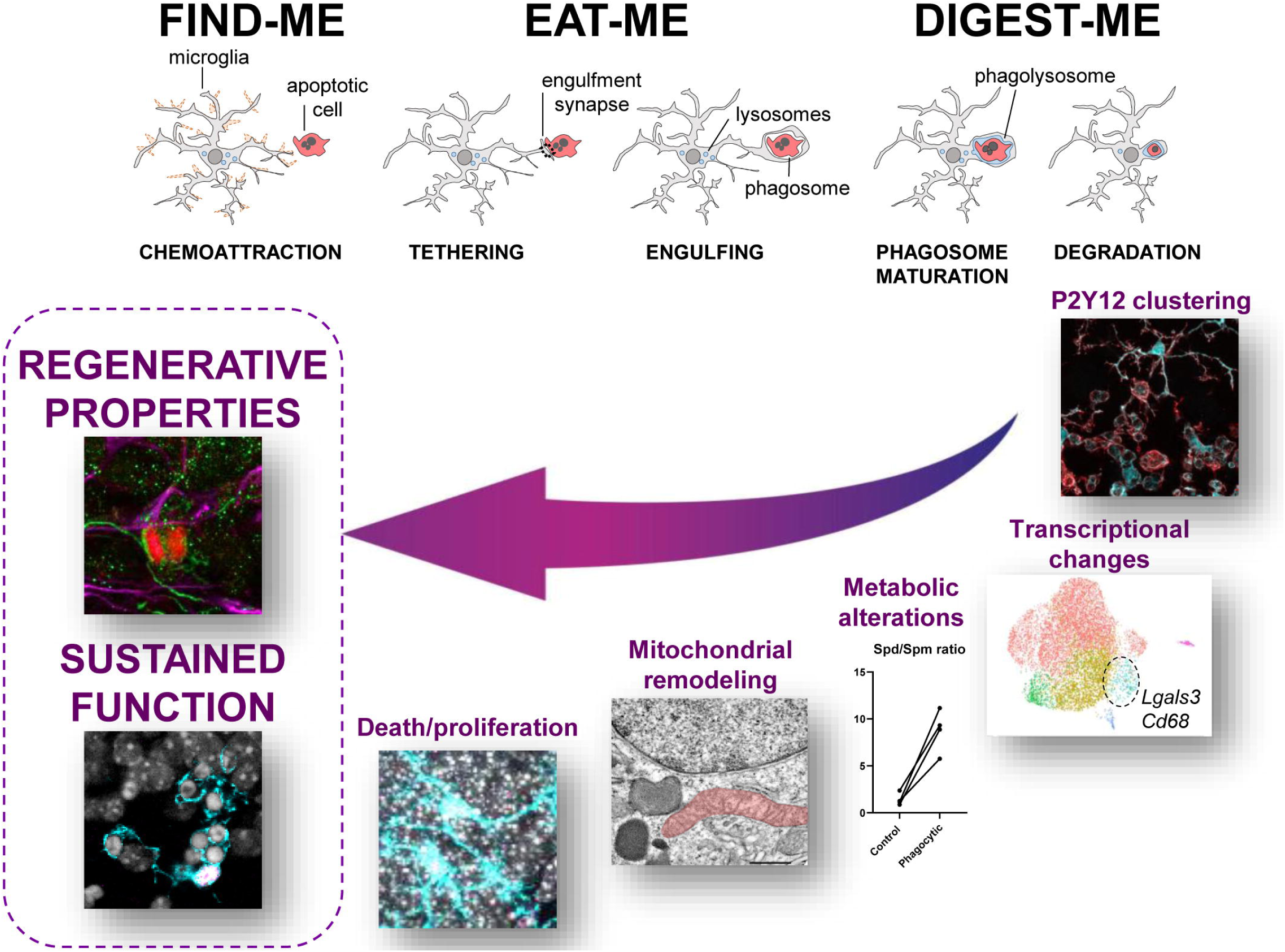
Cartoon summarizing microglial post-phagocytosis adaptations. Here, we show that the canonical stages of phagocytosis (find-me, eat-me, digest-me) are followed by a time-dependent cascade of events, which is initiated by the clustering of P2Y12R on phagocytic pouches. Next, microglia undergo transcriptional changes that include the overexpression of *Cd68* and *Lgals3*; metabolic alterations, including the production of polyamines; mitochondrial remodeling; oxidative stress and cell death; and abortive proliferation. Ultimately, these adaptations allow microglia to maintain their phagocytosis efficiency over time and mediate the regeneration of the neurogenic niche.

### The transcriptional signature(s) associated with microglial phagocytosis

We here identify genes expressed in early post-phagocytic microglia, including lysosomal genes (*Cstb, Cstz, Cd68*) as well as increased expression of *Lgals3*. None of the post-phagocytic microglia genes are included in the Phagocytosis Gene Ontology term (GO:000690), which is severely skewed towards genes involved in the recognition, engulfment, and phagolysosome assembly, i.e., genes that are a pre-requisite for phagocytosis, rather than terms that imply cells actually engaged in phagocytosis.

Phagocytosis has also been linked to other transcriptional signatures, such as DAM, characterized by *Trem2* (triggering receptor expressed in myeloid cells 2) and *Clec7a* (dectin 1)^3^; PAM, which shows some overlap with DAM and is characterized by *Clec7a* and *Gpnmb* (osteoactivin)^4^, and IRM, characterized by *Ifit3* ^5^. The PAM/DAM signature seems to be related to genes required to perform engulfment, as CLEC7a^+^ microglia are more engaged in engulfing than CLEC7a^-^ cells^5^. In contrast, IRM includes genes required to execute degradation, as microglia deficient in the IFN-I receptor *Ifnar1* show a bubbly phenotype with distended lysosomes^5,43^. We did not find changes in expression in the canonical DAM, PAM, and IRM genes *(Trem2*, *Clec7a*, *Gpnmb, Ifit3*) in post-phagocytic microglia. However, some of the post-phagocytic microglia genes (*Lgals3*, *Cd9* or *Lpl)* are upregulated in DAM/PAM, suggesting an overlap between different transcriptional signatures that prevents assigning them unique functional roles.

It would be tempting to brand CD68 and Gal3 as markers of post-phagocytic microglia, but the collective evidence does not suggest that they are uniquely associated with phagocytosis. For example, we here show that the lysosomal CD68 is expressed in many populations of microglia, including non-phagocytic cells and, in the past, we observed increased expression of CD68 during inflammatory conditions^11^. Furthermore, lysosomes are produced and consumed during cell processes unrelated to phagocytosis, including autophagy^44^. Similarly, Gal3 is a pleiotropic sugar-binding protein expressed in several cell types, with both membrane and released forms, and involved in multiple cellular functions, including cell adhesion, proliferation, apoptosis, inflammation and angiogenesis, to name a few^45^. Thus, neither CD68 nor Gal3 should be used as specific markers of phagocytosis.

Therefore, presumed phagocytosis markers, the Phagocytosis GO term, or the DAM/PAM and IRM signatures are not *bona fide* evidence that microglia are engaged in phagocytosis. Instead, our results suggest relying on direct evidences of phagocytosis by quantifying microglial phagocytic pouches, whose identification can be guided by the clustered expression of P2Y12R. Together, the above results suggest that phagocytosis is not a static state, but a time-dependent process encompassing chemotaxis, cargo recognition, engulfment, degradation, and post-phagocytosis adaptations to the metabolic overload. Each stage activates different pathways in microglia and, thus, there is not a unique signature of microglial phagocytosis, but a spectrum of signatures reflecting each of these stages.

### Phagocytosis induces stress and metabolic rewiring

Microglia are considered the “brain professional phagocytes” because of their efficiency in detecting and removing apoptotic cells from the brain parenchyma compared to other cell types^11^. However, this ability does not come without a cost, and here we show that phagocytosis is a stressful process for microglia, which suffer adaptations in response to the metabolic burden imposed by the ingested cargo. These adaptations will likely depend on the nature of the cargo (apoptotic or necrotic debris, β amyloid, α synuclein, synaptic debris, or microbes) as well as the amount ingested, although we did not directly test it. These metabolic adaptations may also be responsible for the abortive proliferation that we found in late post-phagocytic microglia, which we speculate could serve to renovate organelles^46^ damaged from the phagocytosis-induced oxidative stress.

Despite this metabolic stress, post-phagocytic microglia are maintain efficient phagocytosis in the long term when encountering sequential apoptotic challenges, as demonstrated here after LCI and in a model of irradiated glioblastoma, suggesting that post-phagocytosis adaptations are adaptive and constitute part of a recovery program. Similarly, peripheral macrophages also experience adaptations to apoptotic cell phagocytosis that allow them to maintain continued uptake, including increased glycolysis and mitochondrial fission^6,7^. Other macrophages also experience the Warburg effect (dependence on glycolysis instead of oxidative phosphorylation under aerobic conditions)^47^ typical of cancer cells during trained immunity. In this process, an initial stimulus, such as components of the bacterial and fungal wall (β glucan or lipopolysaccharides) triggers epigenetic reprograming and long-term enhanced responses^48^. However, our results do not support that efferocytosis induces trained immunity in microglia: 1, we did not observe a canonical Warburg effect because while post-phagocytic microglia rely more on glycolysis than on oxidative phosphorylation, both pathways are shut down; 2, the transcriptional signature of post-phagocytic microglia is transient, indirectly ruling out major epigenetic changes; and 3, we found no evidences of increased efficiency after sequential apoptotic challenges. Furthermore, we also found increased production of polyamines and other metabolites with antioxidant properties, which post-phagocytic microglia likely use to counteract the metabolic stress associated to the ingested cargo. These findings reveal complex metabolic adaptations to phagocytosis, whose full impact on microglia and neighboring cells could hold therapeutic value in neurodegenerative conditions.

### The regenerative potential of microglial phagocytosis

Our results demonstrate an intricate link between cell death and regeneration provided by post-phagocytic microglia. We tested the functional impact of post-phagocytosis adaptations using Gal3 deficient mice, which had a compromised recovery of rNPC proliferation and newborn neurons after depletion of the neurogenic niche by LCI. While we have not explored the precise mechanism through which Gal3 impacts neurogenesis we speculate that released Gal3, which binds to β-galactoside residues^31^, may directly interact with glycoproteins present in rNPCs and/or neuroblasts. Specifically, beta galactosidase, the enzyme that removes beta galactoside residues, is considered a biomarker of cellular senescence and cell cycle arrest^49^ and it has been shown to promote neuronal differentiation^50^. Another potential mechanism could be mediated by the Gal3 binding protein (Gal3BP), whose release by neural progenitors controls their positioning, possibly by regulating their adhesion to the extracellular matrix^51^. These mechanisms can explain the pro-neurogenic effect of Gal3 derived from post-phagocytic microglia.

The regenerative properties of post-phagocytic microglia revealed in Gal3 deficient mice agree with our previous findings showing that the secretome of phagocytic microglia shapes adult hippocampal neurogenesis by providing a negative feedback loop on newborn neurons^10^. Phagocytic microglia increase the expression of growth factors, matrix remodeling proteins, and cytokines, all of which have been involved in neurogenesis, although the specific molecule/s responsible were not identified^10^. We speculate that metabolites released by post-phagocytic microglia may also contribute to the regulation of the neurogenic niche. For instance, spermidine acts on several stem cell types^52–54^ and, as an autophagy inducer^55^, it also has a strong potential to regulate neurogenesis^56^ that needs to be experimentally validated.

However, the regenerative properties of post-phagocytic microglia may be useful in neurodegenerative diseases but detrimental in other pathological states, such as brain cancer. Gliomas are the most common type of primary brain tumors, among which, glioblastomas are the most aggressive. Brain tumors are usually first treated surgically to remove the tumor mass, then with radiotherapy and chemotherapy to kill the remaining tumor cells^57^. Dead tumor cells are phagocytosed by TAMs, as we show here, enhancing their immunosuppressive state^58^. Our results suggest that post-phagocytosis adaptations in TAMs may directly impact tumors, possibly through metabolites like spermidine, which has been recently shown to promote tumorigenesis^59^. Like the Yin and Yang, phagocytosis adaptations may be advantageous for brain regeneration but harmful in cancer.

In summary, we have identified the complex set of transcriptional and metabolic alterations endured by post-phagocytic microglia, which allow them to maintain continued phagocytosis efficiency and promote the recovery of the neurogenic niche.

## ACKNOWLEDGEMENTS

This work was supported by grants from the Spanish Ministry of Science and Innovation Competitiveness MICIU/AEI/10.13039/501100011033 and by “ERDF A way of making Europe” (RTI2018-099267-B-I00, PID2022-136698OB-I00 and RYC-2013-12817 to AS, and RTI2018-102260-B-I00, PID2021-129053OB-I00 to JPLA); by Basque Government grants (IT1473-22 and PIBA 2020_1_0030), a Tatiana Foundation Award (P-048-FTPGB 2018), and an Alzheimer Association award (AARG-NTF-24-1304352) to AS. It was also supported by Spanish Ministry of Science and Innovation Competitiveness (PID2022-140525NB-I00) and the Excellence Unit IBRAINS-IN-CYL, Strategic plans and strategic research programmes of excellence from the Regional Government of Castilla y León, co-funded by the ERDF Operational Programme (CLU-2023-1-01) to JV; by the Basque Government Elkartek Program (BG24), Spanish Ministry of Science and Innovation Competitiveness (PID2021-125104OB-I00), Fundación Científica de la Asociación Española Contra el Cáncer (AECC PRYGN247108ASPI) and Alzheimeŕs Association EV-METABOLIC Project (AARG-NTF-22-968911) to JMFP; and by an ISCIII-FEDER PI20/01063 to MA; and by Generalitat Valenciana (CIPROM/2023/15) to JPLA. This work was also supported by the Spanish Ministry of Science, Innovation and Universities/AEI (PID2024-157400OB-I00) to JLV. MMR holds a University of the Basque Country EHU predoctoral fellowship and XCP a Basque Government predoctoral fellowship, and MGD, MPI and LA are recipients of predoctoral fellowships from the Spanish Ministry of Science and Innovation. SB is a recipient of a predoctoral fellowship from the Spanish Ministry of Economy and Competitiveness. The Institute of Neurosciences is a Severo Ochoa Center of Excellence (grant CEX2021-001165-S, funded by MCIU/AEI/10.13039/501100011033). We are grateful for the technical support of Victor Sánchez-Zafra and Jikke Wagendorp. Achucarro and UPV/EHU SGIker technical and human support is gratefully acknowledged.

---

**Supp. Figure 1.**
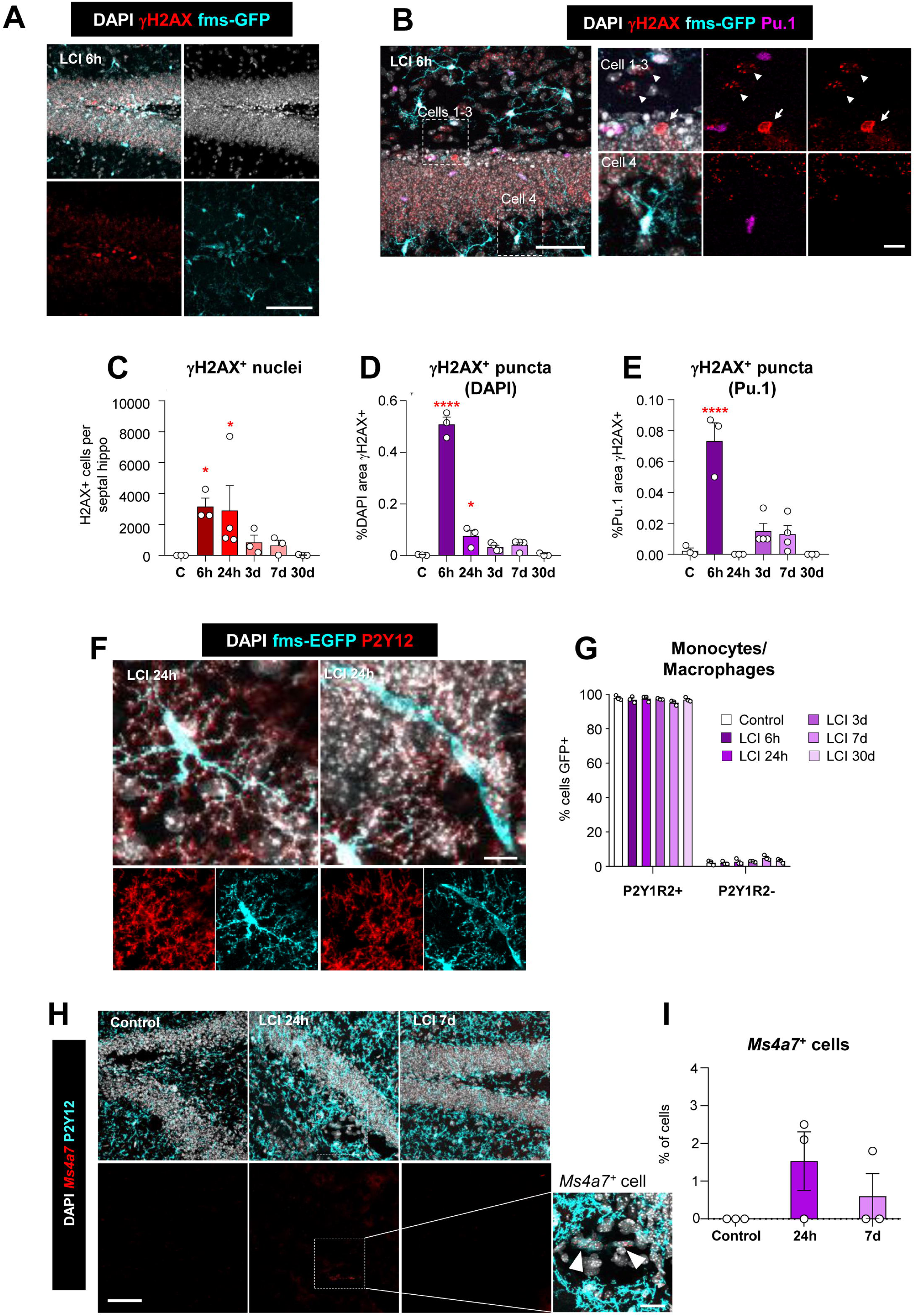
LCI results in minimal DNA damage in microglia and monocyte infiltration. **(A)** Representative confocal images of double-strand breaks labeled with γH2AX at 6h after LCI showing DAPI (white), γH2AX (red) and fms-EGFP (cyan) staining 6h after LCI. **(B)** Representative images of a γH2AX^+^ nucleus (red) indicated with a white arrow and other nuclei labeled with γH2AX puncta indicated by white arrowheads (cells 1-3). A microglial nucleus labeled with DAPI (white), Pu.1 (magenta), and fms-EGFP (cyan) not expressing γH2AX is shown (cell 4). **(C)** Number of γH2AX^+^ nuclei in the DG per septal hippocampus. **(D)** Percentage of nuclear volume (DAPI) occupied by γH2AX^+^ puncta. **(E)** Percentage of microglial nuclear volume (Pu.1) occupied by γH2AX^+^ puncta. **(F)** Representative confocal images of microglia and monocytes/macrophages at 24h after LCI. Microglia are labeled with both P2Y12 (red) and fms-EGFP (cyan), while monocytes/macrophages only express fms-EGFP (cyan). DAPI (white) staining shows nuclei. The elongated, smooth-surfaced fms-EGFP^+^, P2Y12-cell is likely a perivascular macrophage. **(G)** Percentage of fms-EGFP^+^ cells identified as microglia (P2Y12^+^) and monocyte/macrophages (P2Y12^-^) **(H)** Representative images of the monocytes/macrophage gene *Ms4a7* by RNAScope, showing DAPI (white), *Ms4a7* mRNA (red) and P2Y12 (cyan) staining in the DG 24h and 7d after LCI. The white dotted rectangle shows in higher magnification two *Ms4a7+* cells (white arrowheads) that are not microglia (P2Y12^-^) **(I)** Percentage of *Ms4a7^+^* cells in the DG 24h and 7d after LCI. Bars show mean ± SEM of n=3-4 mice. Data was analyzed by one-way ANOVA followed by Holm-Sidak post hoc tests when appropriate. When homoscedasticity was not accomplished a Kruskal-Wallis test was performed [C]. Asterisks represent significance between control and LCI (6h, 24h, 3d, 7d, 30d). * represents p<0.05, *** represents p<0.001. Scale bars= 100μm [A, B, H], 10μm (inserts in [B]); 10μm [F]. z=16.8μm [A]; z=26.6μm, z=0.7μm [B]; z= 17.5μm [F]; z=16.45μm, z=17.5, μm z=8.75μm [D]; [H, left to right];

**Supp. Figure 2.**
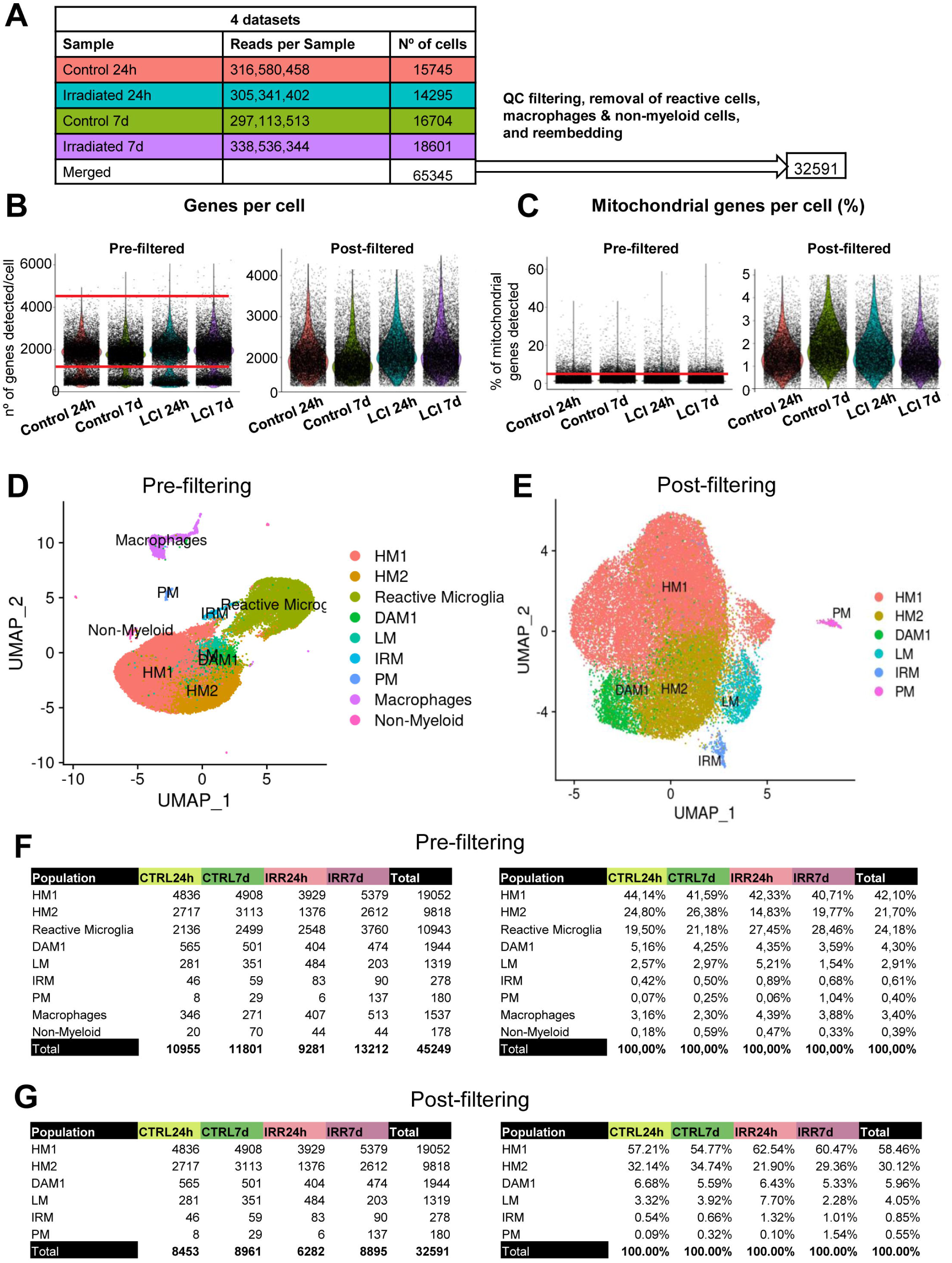
Single-cell RNA-seq processing of microglia after LCI. **(A)** Summary table showing the total number of sequenced cells per condition and the final cell count following quality control filtering and microglia subset selection **(B)** Violin plots depicting the distribution of genes counts detected per cell before (left) and after (right) quality control filtering. Red lines denote the filtering. **(C)** Violin plots illustrating the percentage of mitochondrial genes detected per cell before (left) and after (right) quality control filtering. Red lines indicate the filtering thresholds. **(D)** UMAP visualization of the supervised clustering annotation, based on the 30 most significant principal components (PCs) derived from Principal Component Analysis (PCA). **(E)** UMAP representation of selected clusters, excluding non-myeloid cells, macrophages, and ex vivo reactive microglia. **(F)** Summary tables reporting the number (left) and percentage (right) of cells detected per population and treatment condition shown in D. **(G)** Summary tables reporting the number (left) and percentage (right) of cells detected per microglia subpopulation and treatment condition shown in E.

**Supp. Figure 3.**
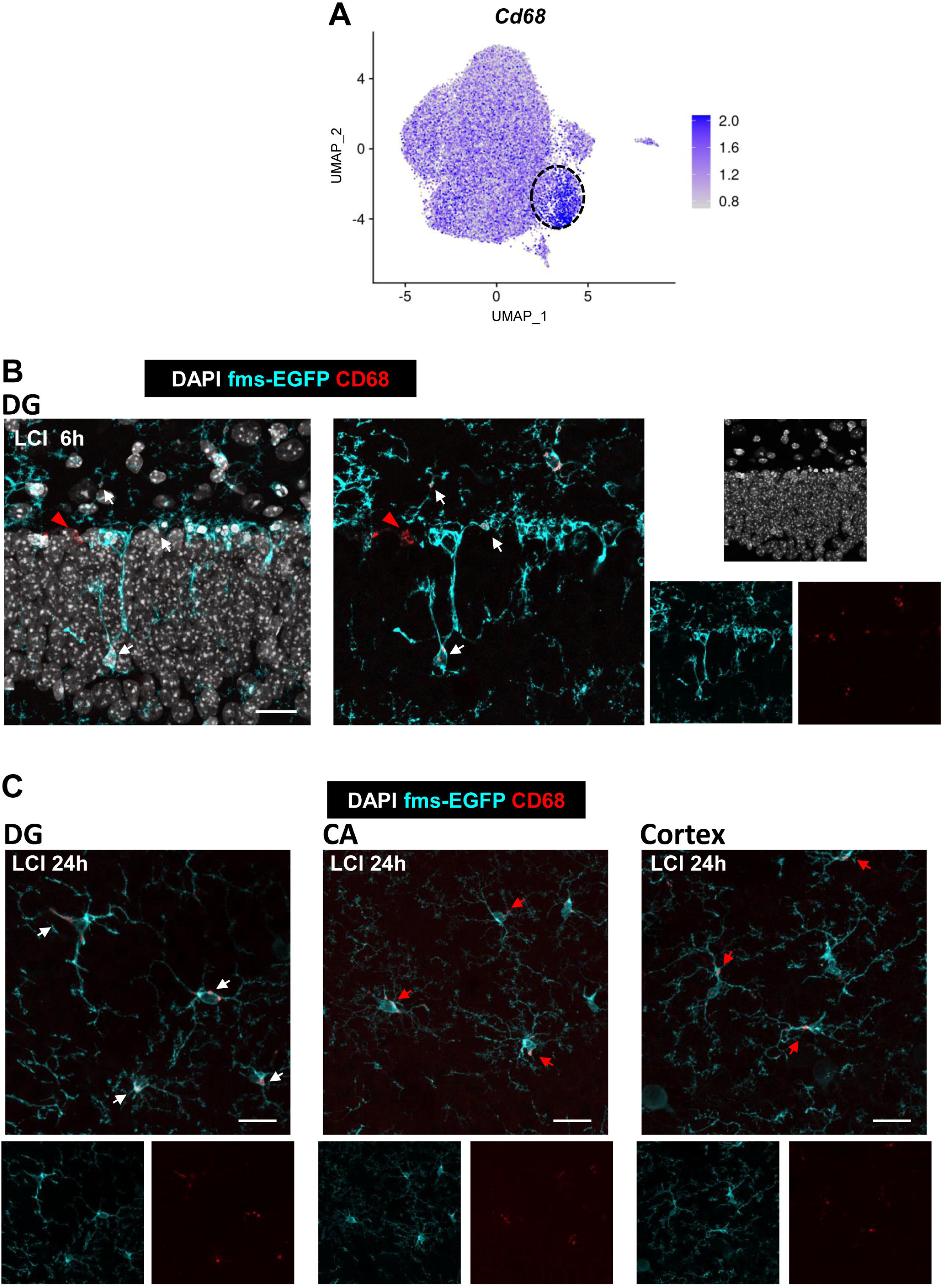
CD68 is not a specific marker of post-phagocytic microglia. **(A)** UMAP showing *Cd68* gene expression after LCI. The LM cluster is highlighted by a dotted black circle. **(B)** Representative confocal image of CD68^+^ puncta in phagocytic microglia (white arrows) and non-microglial cells (red arrowhead) 6h after LCI, showing DAPI (white), CD68 (red), and fms-EGFP (cyan). **(C)** Representative confocal images of CD68^+^ post-phagocytic microglia from the DG (white arrows) and non-phagocytic microglia (red arrows) from the CA and the cortex 24h after LCI, showing DAPI (white), CD68 (red), and fms-EGFP (cyan). Scale bars= 20μm [B, C]. z=16.8μm [B]; z=11.2μm, z=16.8μm, z=19.6μm [C, left to right].

**Supp. Figure 4.**
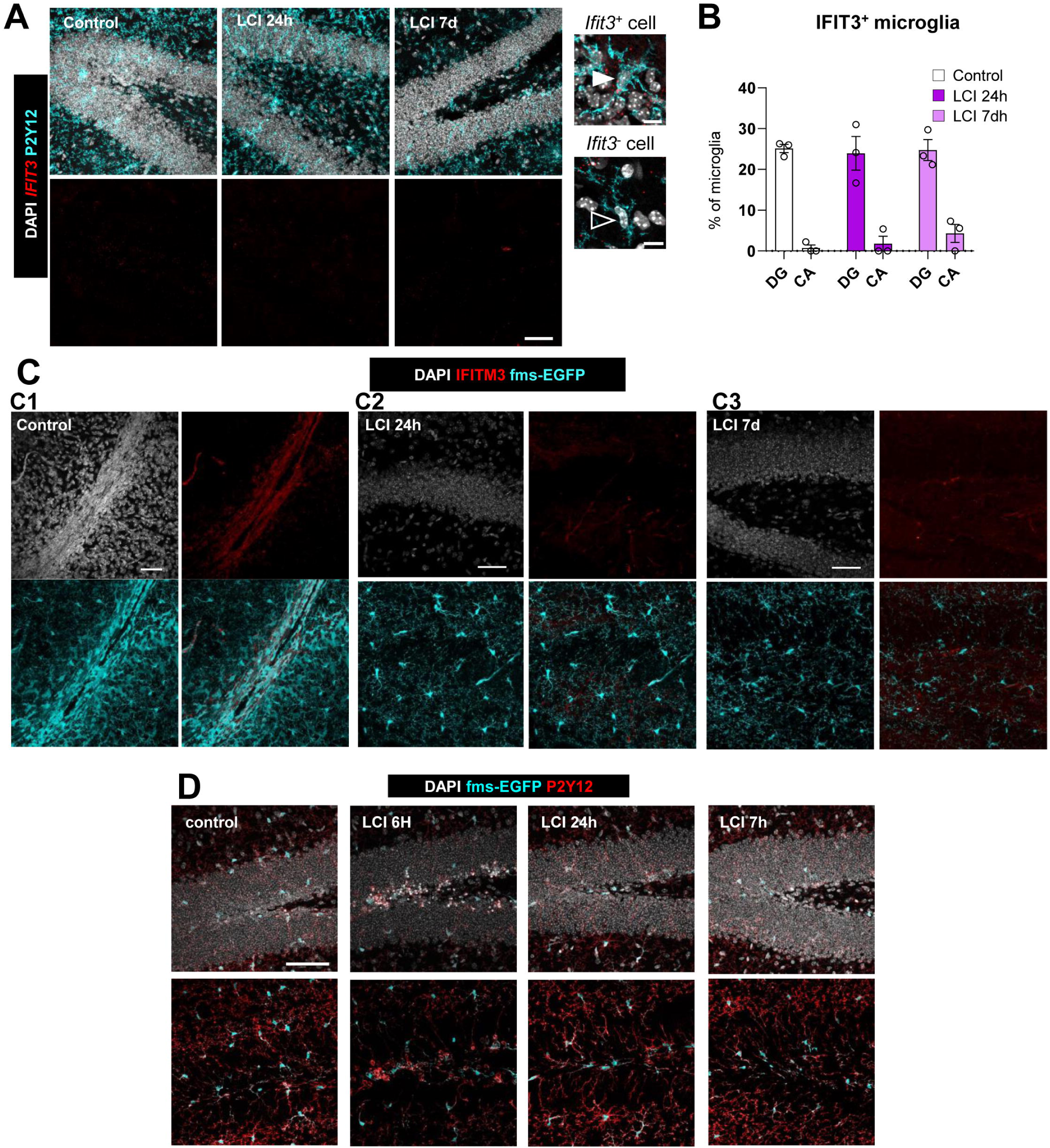
The IRM cluster does not contain post-phagocytic microglia. **(A)** Representative confocal images of the IRM signature gene *Ifit3* by RNAScope, showing DAPI (white), *Ifit3* mRNA (red) and P2Y12 (cyan) staining in the DG 24h and 7d after LCI. The white arrowhead points to an *Ifit* ^+^ microglia, and the black arrowhead to an *Ifit3* ^-^ microglia. **(B)** Analysis of the percentage of *Ifit3* ^+^ microglia in the DG and CA 24h and 7d after LCI. **(C)** Representative images of the IRM signature protein IFITM3 showing DAPI (white), IFITM3 (red) and P2Y12 (cyan) staining at 24h and 7d after LCI. The control image shows a large blood vessel labeled with IFITM3, whereas no staining was detected in microglia from the DG 24h and 7d after LCI. **(D)** Representative confocal images of P2Y12 expression in the DG after LCI, showing DAPI (white), P2Y12 (red) and fms-EGFP (cyan) staining. Bars show mean ± SEM of n=3 mice. Data was analyzed by one-way ANOVA followed by Holm-Sidak post hoc tests when appropriate. Scale bars=50μm [A,C, D], 10μm (inserts in [A,D]); z=12.25μm, z=11.2μm, z=10.15μm [A]; z=32.9μm [C1], z=27.3μm [C2], z=24.5μm [C3]; z=16.1μm, z=18.9μm, z=20.3μm, z=12.6μm [D, left to right].

**Sup Fig. 5.**
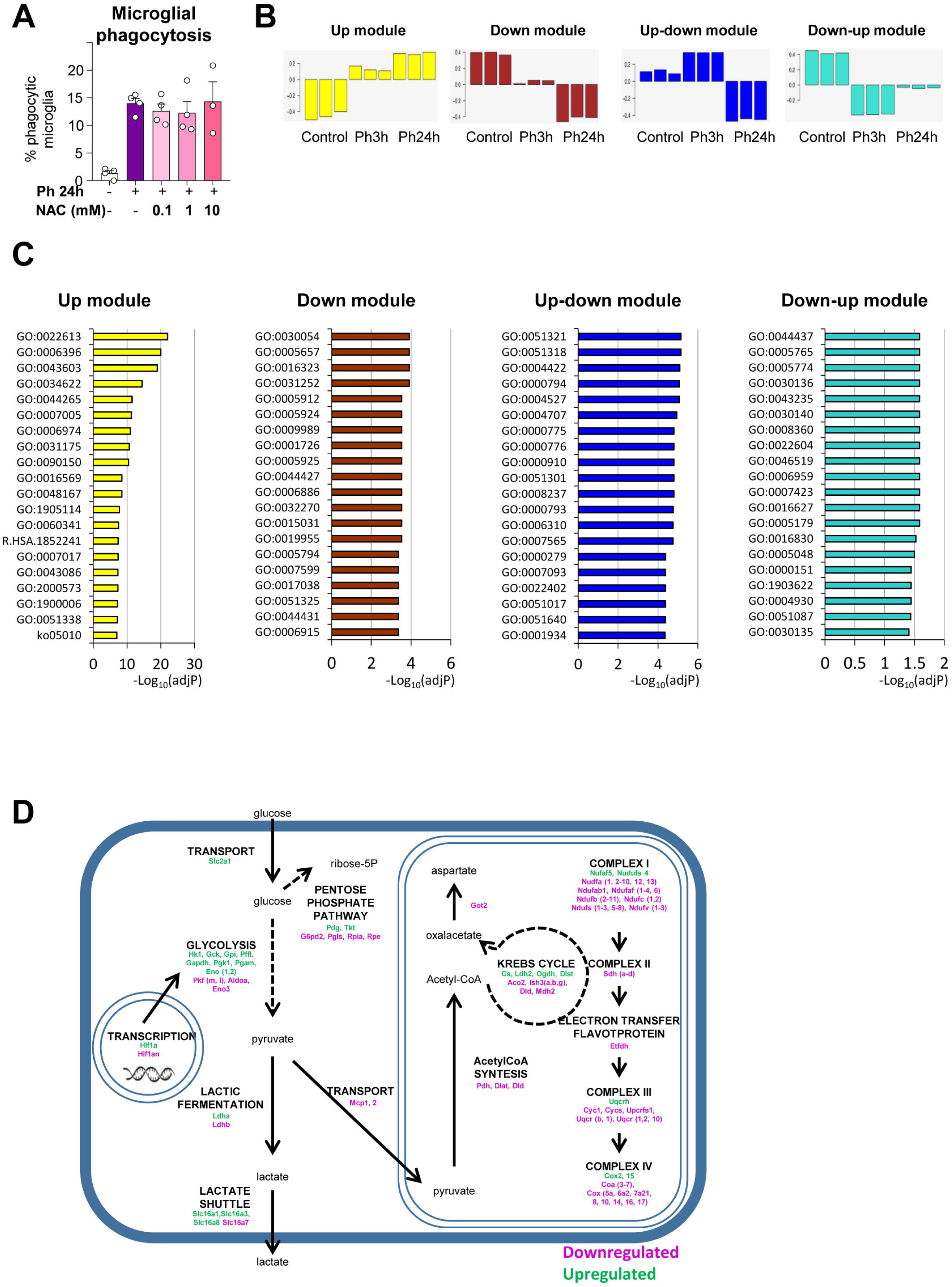
Functional analysis of phagocytic microglia by WGCNA and IPA analysis. **(A)** Percentage of phagocytic microglia after phagocytosis *in vitro* (24h) and pre-treatment with the antioxidant N-Acetylcysteine (NAC). **(B)** Modules obtained by WGCNA analysis of gene arrays comparing primary cultures of naïve and phagocytic microglia at 3h and 24h. **(C)** IPA analysis showing the top 20 significantly changed GO terms associated with each module. Up module (yellow): GO:0022613, ribonucleoprotein complex biogenesis; GO:0006396, RNA processing; GO:0043603, cellular amide metabolic process; GO:0034622, cellular macromolecular complex assembly; GO:0044265, cellular macromolecule catabolic process; GO:0007005, mitochondrion organization; GO:0006974, cellular response to DNA damage stimulus; GO:0031175, neuron projection development; GO:0090150, establishment of protein localization to membrane; GO:0016569, covalent chromatin modification; GO:0048167, regulation of synaptic plasticity; GO:1905114, cell surface receptor signaling pathway involved in cell-cell signaling; GO:0060341, regulation of cellular localization; R.HSA.1852241, organelle biogenesis and maintenance; GO:0007017, microtubule-based process; GO:0043086, negative regulation of catalytic activity; GO:2000573, positive regulation of DNA biosynthetic process; GO:1900006, positive regulation of dendrite development; GO:0051338, regulation of transferase activity; ko05010, Alzheimer’s disease. Down module (brown): GO:0030054, cell junction; GO:0005657, replication fork; GO:0016323, basolateral plasma membrane; GO:0031252, leading edge; GO:0005912, adherens junction; GO:0005924, cell substrate adherens junction; GO:0009989, cell matrix junction; GO:0001726, ruffle; GO:0005925, focal adhesion; GO:0044427, chromosomal part; GO:0006886, intracellular protein transport; GO:0032270, positive regulation of cellular protein metabolic process; GO:0015031, protein transport; GO:0019955, cytokine binding; GO:0005794, Golgi apparatus; GO:0007599, hemostasis; GO:0017038, protein import; GO:0051325, interphase; GO:0044431, golgi apparatus part; GO:0006915, apoptotic program. Up-Down (blue) module: GO:0051321, meiotic cell cycle; GO:0051318, G1 phase; GO:0004422, metalloendopeptidase activity; GO:0000794, condensed nuclear chromosome; GO:0004527, exonuclease activity; GO:0004707, MAP kinase activity; GO:0000775, chromosome centromeric region; GO:0000776, kinetochore; GO:0000910, cytokinesis; GO:0051301, cell division; GO:0008237, metallopeptidase activity; GO:0000793, condensed chromosome; GO:0006310, DNA recombination; GO:0007565, female pregnancy; GO:0000279, M phase; GO:0007093, mitotic cell cycle checkpoint; GO:0022402, cell cycle process; GO:0051017, actin filament bundle formation; GO:0051640, organelle localization; GO:0001934, positive regulation of protein amino acid phosphorylation. Down-Up module (turquoise): GO:0044437, vacuolar part; GO:0005765, lysosomal membrane; GO:0005774, vacuolar membrane; GO:0030136, clathrin coated vesicle; GO:0043235, receptor complex; GO:0030140, trans Golgi network transport vesicle; GO:0008360, regulation of cell shape; GO:0022604, regulation of cell morphogenesis; GO:0046519, sphingoid metabolic process; GO:0006959, humoral immune response; GO:0007423, sensory organ development; GO:0016627, oxidoreductase activity acting on the ch-ch group of donors; GO:0005179, hormone activity; GO:0016830, carbon carbon lyase activity; GO:0005048, signal sequence binding; GO:0000151, ubiquitin ligase complex; GO:1903622, transcription from RNA polymerase III promoter; GO:0004930, G protein coupled receptor activity; GO:0051087, chaperone binding; GO:0030135, coated vesicle. **[D]** Cartoon representing genes codifying for metabolic enzymes upregulated (green) and downregulated (magenta) 24h after phagocytosis in microglia. Raw data was published in ^10^.

**Supp. Figure 6.**
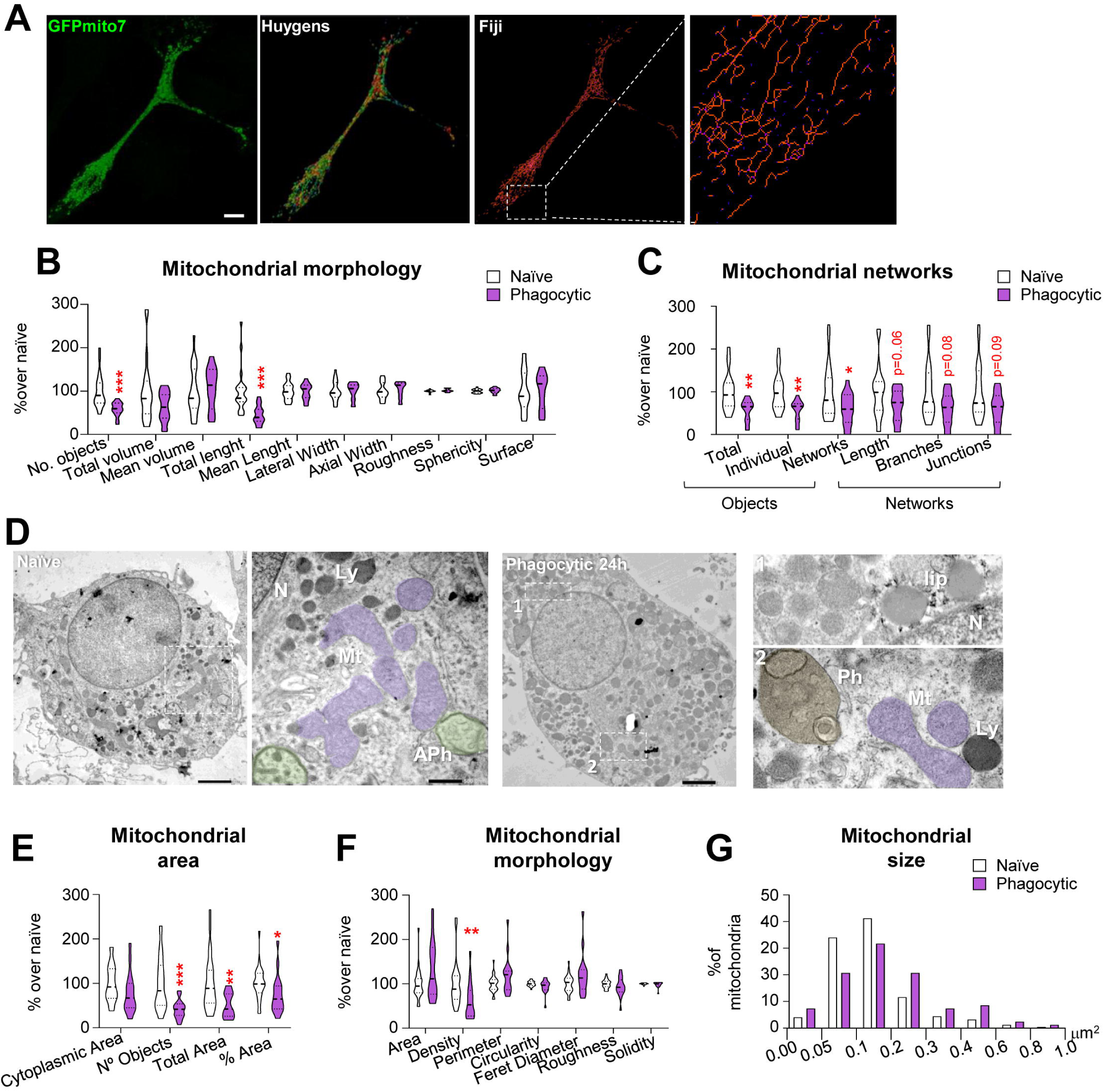
Phagocytosis remodels the microglial mitochondrial network. **(A)** Comparison of the methods used to assess mitochondria morphology in naïve and phagocytic BV2 microglia using confocal images Mitochondria were labelled with GFPmito7 (green) and mitochondrial morphology and networks were assessed by Huygens and Fiji, respectively. **(B)** Analysis of mitochondrial morphology by Huygens showing the percentage of different mitochondrial parameters in BV2 naïve and phagocytic microglia. **(C)** Analysis of mitochondrial networks by Fiji showing the percentage of different mitochondrial parameters in BV2 naïve and phagocytic microglia. **(D)** Representative TEM images of primary naïve and phagocytic microglia showing mitochondria (pink,Mt) and other organelles, such as lysosomes (dark,Ly). In control cells, abundant primary and secondary autophagosmes (purple and blue, respectively; APh) were found, whereas in phagocytic cells several degradative organelles such as multilamelated bodies (green,MLB) were found. Lipidic content (lip) and phagosomes (Ph) were also found in phagocytic cells. **(E)** TEM analysis of mitochondrial object number and area per microglial cytoplasmic area in microglial cytoplasm in primary naïve and phagocytic microglia. **(F)** TEM analysis of mitochondrial morphology in primary naïve and phagocytic microglia. **(G)** Histogram of mitochondrial objects according to their size (% of mitochondria) in primary naïve and phagocytic microglia. Data shows the information of n=3 independent experiments. Violin plots show the data distribution including extreme values; lower and upper hinges correspond to the third quartile respectively. Data was analyzed by Student’s t-test. Some data [B,C,E,F] were log-transformed to comply with homoscedasticity and/or normality. * indicates p <0.05, ** indicates p <0.01, *** indicates p <0.001. Scale bars= 10μm [A], 2m for whole cell images and 0.5μm for details [D]. z=7.64μm [A].

**Supp Fig. 7.**
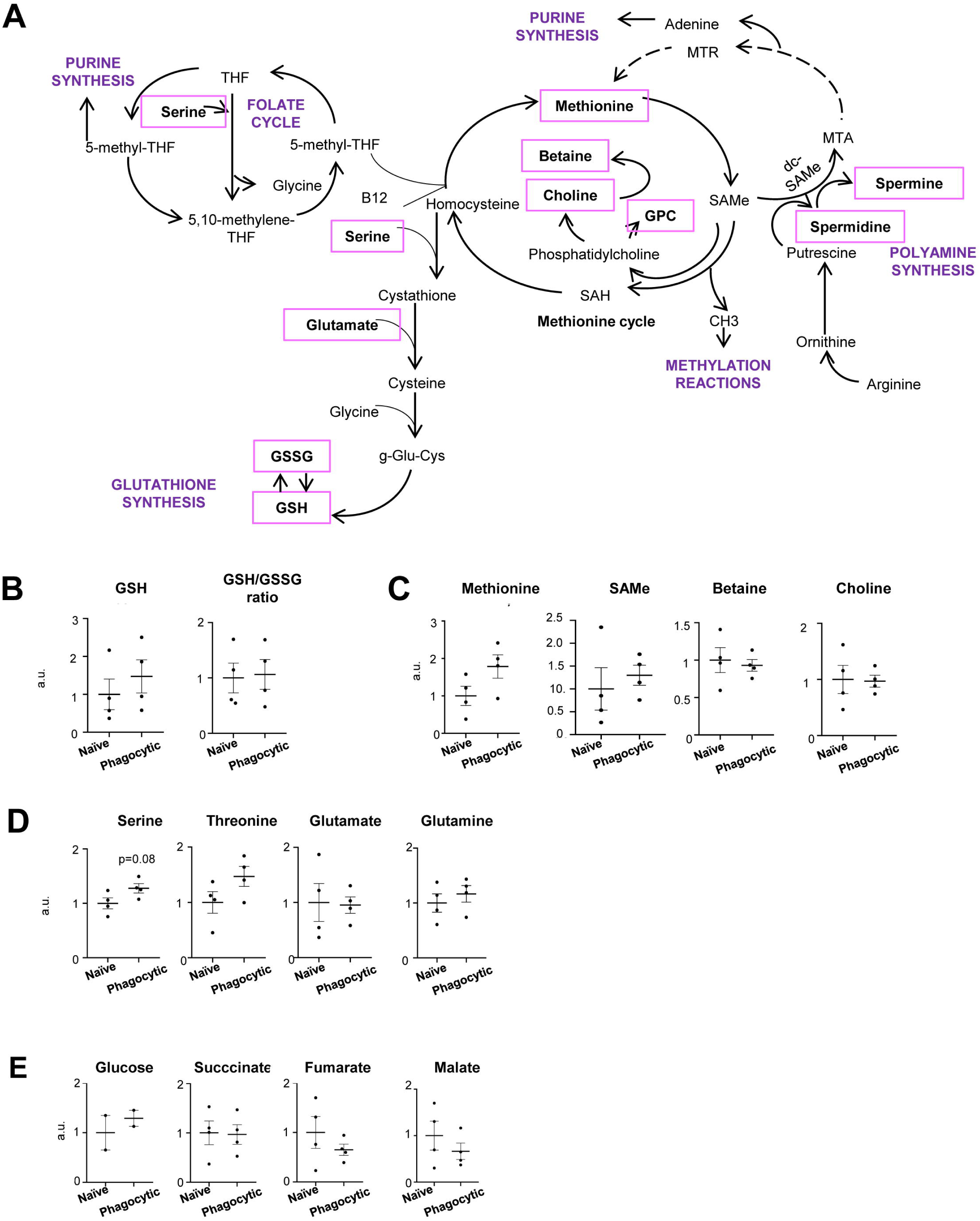
Phagocytosis alters the synthesis pathways of polyamines and glutathione in microglia. **(A)** Graphical cartoon of the analyzed metabolites (purple) by mass spectrometry and the metabolic pathways in which they are involved. **(B)** Relative production of GSH in arbitrary units (a.u.) and GSH/GSSG ratio in naïve and phagocytic microglia. **(C)** Relative production of methionine, SAMe, betaine and choline in arbitrary units (a.u.) in naïve and phagocytic microglia. **(D)** Relative production of serine, threonine, glutamate and glutamine in arbitrary units (a.u.) in naïve and phagocytic microglia. **(E)** Relative production of glucose, succinate, fumarate, and malate in arbitrary units (a.u.) in naïve and phagocytic microglia.

**Supp. Fig. 8.**
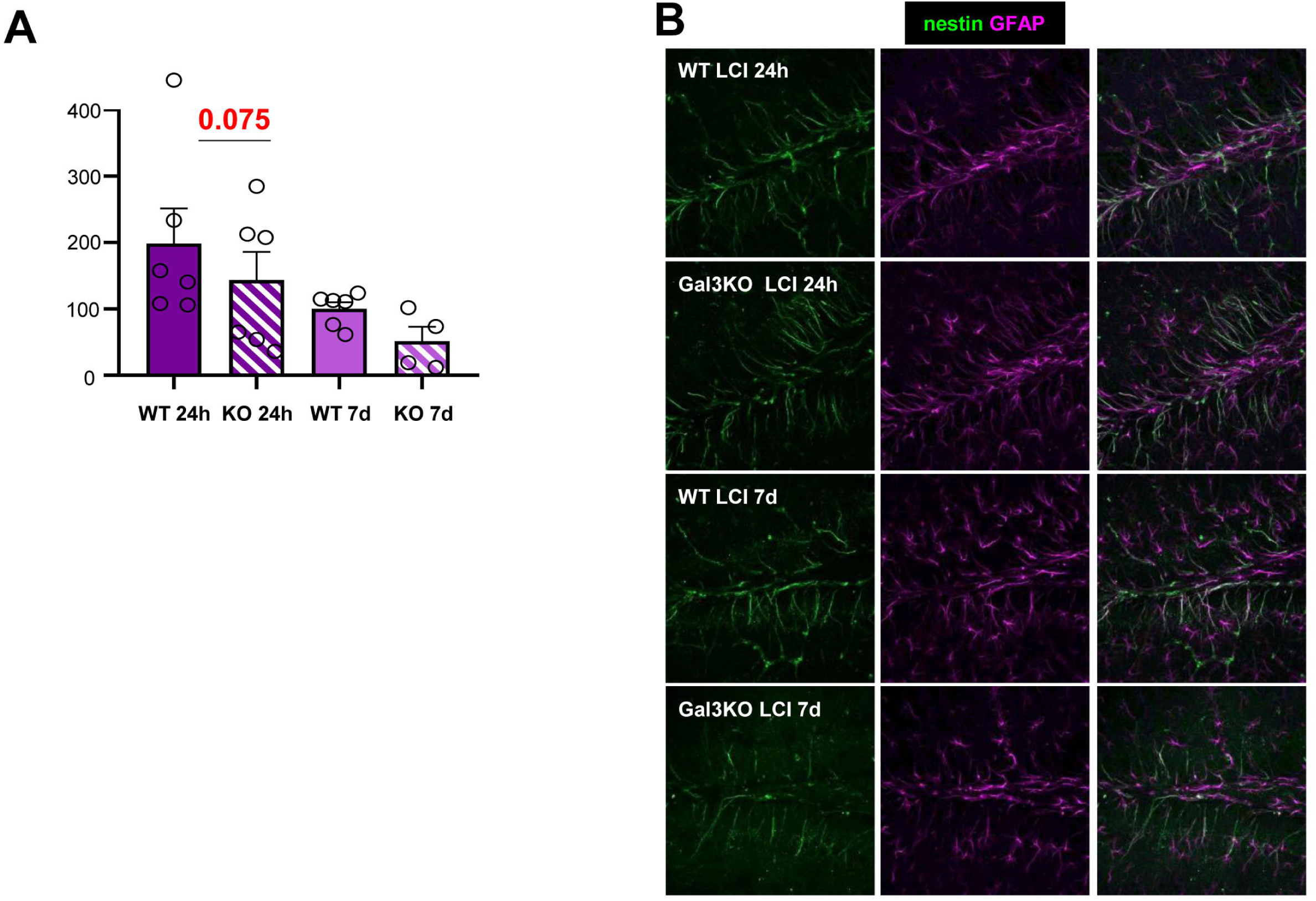
Galectin 3 deficient mice fail to recover the neurogenic niche after LCI. **(A)** Relative density of mature (CD, EF) neuroblasts in the DG per septal hippocampus. **(B)** Representative image of the hippocampus showing DAPI (white), the proliferation marker Ki67 (red), and the radial processes of the rNPCs labeled with both nestin (green) and GFAP (magenta). Bars show the mean ± SEM of n=4-6 mice. Neuroblasts were analyzed as relative densities (compared to the WT group 7 days after LCI) because this experiment was analyzed in 2 independent batches of animals, each comprising the four experimental groups. Data was analyzed by one-way ANOVA. Scale bars=100μm. z=16.8, 20.3, 16.8, 16.8 μm (top to bottom).

